# Wide visual angle enhances social interaction in a realistic virtual reality environment under simulated artificial vision

**DOI:** 10.1101/2025.03.27.645647

**Authors:** Sandrine Hinrichs, Yvain Tisserand, Elsa Scialom, Diego Ghezzi

**Affiliations:** Laboratory of Psychophysics, Brain Mind Institute, School of Life Sciences, École Polytechnique Fédérale de Lausanne, Lausanne, Switzerland; Ophthalmic and Neural Technologies Laboratory, Department of Ophthalmology, University of Lausanne, Hôpital ophtalmique Jules-Gonin, Fondation Asile des Aveugles, Lausanne, Switzerland; Swiss Center for Affective Sciences, University of Geneva, Geneva, Switzerland

**Keywords:** simulated artificial vision, visual field size, virtual reality, retinal implants, social interaction

## Abstract

*Objective*. Social interaction largely relies on visual perception. Yet artificial vision research has primarily focused on navigation and object recognition while assessment of social interaction remains scarce. This study investigates how the visual field size in simulated artificial vision impacts social perception in a virtual reality setting. *Approach*. We assessed the impact of visual field size in sighted participants experiencing simulated artificial vision in a medical practice across three social task categories: indoor identification (searching for an empty chair, identification of body orientation, body appearance, facial expressions, relationship type, role and interpretation of body language), locomotion (tracking and following a moving person), and outdoor observation (person recognition in a cluttered outdoor environment). *Main Results*. A wide range of social perception and locomotion tasks in structured indoor environments can be completed with a 45° field of view in simulated artificial vision. Compared to 20°, the broader visual field not only enhances accuracy but also enables faster task execution and a more precise assessment of one’s own performance. Although the benefits of an expanded visual field are limited in cluttered environments, a 45° visual field represents a clear advantage in social contexts. *Significance*. These findings highlight the importance of an expanded visual field for artificial vision users in social settings, reinforcing its role in enhancing social cue processing and the ability to follow people. By integrating simulated artificial vision within an immersive virtual reality framework, this study bridges the gap between controlled experimental paradigms and real-world needs and challenges.

## 1. INTRODUCTION

Social interaction is a fundamental component of daily life, enabling individuals to navigate professional, personal, and public environments effectively. It relies on a broad set of perceptual and cognitive skills, including the ability to recognize individuals, interpret body orientation, track motion, identify gestures and facial expressions, and analyze social roles and relationships [1]. Among the sensory modalities contributing to social perception, vision plays a dominant role in providing rapid, high-resolution information about people, their movements, and their intentions [2–4]. Visual cues are essential for interpreting nonverbal communication, coordinating social interactions, and maintaining situational awareness in dynamic environments [5]. Conversely, visual impairment profoundly disrupts social functioning. Individuals with severe vision loss experience increased social isolation and reduced quality of life due to inability to recognize faces, detect emotions, follow conversational cues, and maintain fluid interactions [6, 7]. Impaired vision also hinders spatial awareness and mobility in crowded environments, exacerbating challenges in social engagement [8, 9]. As such, individuals with profound visual impairments significantly rely on compensatory auditory and tactile strategies, though these are insufficient to fully replace the richness and immediacy of visual information [10].

Restorative approaches such as artificial vision through neurostimulation, including visual prostheses and optogenetics, seek to reintroduce visual information to blind people [11–14]. Evaluating the functional impact of these technologies in real-world scenarios remains challenging, as clinical trials are constrained by environmental complexity and limited patient availability. A practical way to assess artificial vision before clinical implementation is through Simulated Artificial Vision (SAV), which allows researchers to systematically investigate perceptual parameters under controlled yet ecologically valid conditions. While SAV has been primarily used to investigate object recognition, navigation, and spatial processing, studies in social environments remain scarce and are often limited to controlled settings that do not reflect the complexities of real-world social interactions [15–18].

In previous work, we modeled phosphene perception from the POLYRETINA device [19, 20], combining SAV with a head-mounted display (HMD) for augmented reality (AR) and behavioral assessments in an artificial street laboratory [21]. Our findings showed that a 45° visual angle significantly improved participants’ ability to navigate and complete common outdoor tasks more effectively, safely, and efficiently than a 20° angle, while further increasing the visual field size provided no additional benefits. Building on this approach, the present study applies the same SAV simulation of 20° and 45° visual angles in virtual reality (VR) to examine how visual field size impacts human behavior in daily social interaction tasks within a virtual medical practice environment. Specifically, this study examines three key functional aspects of artificial vision for social interaction. First, indoor identification tasks assess participants’ ability to recognize body orientation, physical characteristics, facial expressions, and social roles. Second, the locomotion task evaluates their capacity to track and follow a moving individual, requiring continuous spatial awareness and dynamic visual processing. Finally, the outdoor observation task measures participants’ ability to identify and interpret multiple individuals in a cluttered outdoor setting. Our findings reinforce the importance of visual field size in artificial vision for tasks requiring social perception. A wider visual angle facilitates faster, more confident and accurate social perception in indoor identification tasks as well as a better ability to track and follow movements of others. Performance in the outdoor scene suggests that the wide field advantage is diminished in cluttered environments. Understanding these effects is crucial for optimizing prosthetic vision technologies. By integrating SAV in a VR-based naturalistic social environment, this study further contributes in bridging the gap between controlled experimental paradigms and real-world applications in artificial vision research.

## 2. METHODS

### 2.1 Participants

The study is a between-subjects design with participants recruited from students at École Polytechnique Fédérale de Lausanne and Université de Lausanne (**Table 1**). Participants were randomly assigned to one of two viewing condition groups, where they experienced the VR scene in one of two SAV visual fields: 20° circular (SAV 20°) or 45° circular (SAV 45°). The sample size was determined via power analysis (GPower version 3.1.9.7) [22], based on the effect size (d = 1.08) observed in a previous study [21]. 30 participants (15 per group) were required for a two-tailed independent samples t-test with an alpha level of 0.05 and a power of 0.8. This sample size is consistent with prior studies on orientation and mobility in low or artificial vision [23]. Inclusion criteria required participants to be aged between 18 and 45 years, with normal or corrected-to-normal vision, the ability to understand written and spoken English or French, no physical mobility limitations, no neurological disorders or claustrophobia, and no prior experience with SAV. Participation was voluntary, with a financial compensation of 25 CHF per hour. The study was conducted in either English or French, depending on the participant’s preference. Participants were naive to the research question.

**Table 1.** Demographic data for study participants.

|  | SAV 20° | SAV 45° |
| --- | --- | --- |
| **Number** | 15 | 15 |
| **Age (mean ± SD) in years** | 22.93 ± 1.62 | 23.13 ± 4.16 |
| **Age (min - max) in years** | 20 - 26 | 19 - 37 |
| **Sex** | 10 female / 5 male | 9 female / 6 male |
| **Dominant eye** | 12 right / 3 left | 9 right / 6 left |
| **Language** | 9 french / 6 english | 10 french / 5 english |

### 2.2 Simulated artificial vision

Participants wore an HTC VIVE Pro Eye HMD operated using an HP Victus 15L computer equipped with an Intel Core i7 12^th^ generation CPU and a GeForce RTX 3060 Ti GPU. SAV was implemented using Unity (version 2019.2.16f1) and Cg shaders, which transformed the VR space into a phosphene representation (**Figure 1**). Phosphenes are bright spots perceived when visual implants stimulate neurons, forming the basic units of artificial vision [24]. Parameters matched those used in a previous study [21]. Phosphenes were circular, with a base size of 80 µm and a spacing of 120 µm, corresponding to approximately 0.275° and 0.414°, respectively. These parameters were based on the POLYRETINA prosthetic device [16, 25, 26]. A distortion effect mimicking unintended axon fiber activation was set to λ = 2 [16, 26–28]. SAV operated at 5 Hz with an 11 ms frame duration (one frame on, 17 frames off) and was presented exclusively to the dominant eye. Random variability was introduced at each frame: phosphene size varied within ±30% of their base size, brightness varied between 50% (gray) and 100% (white), and 10% of phosphenes were randomly omitted. SAV also accounted for perceptual fading, with a compensation strategy applied as previously described [16, 29].

**Figure 1.**
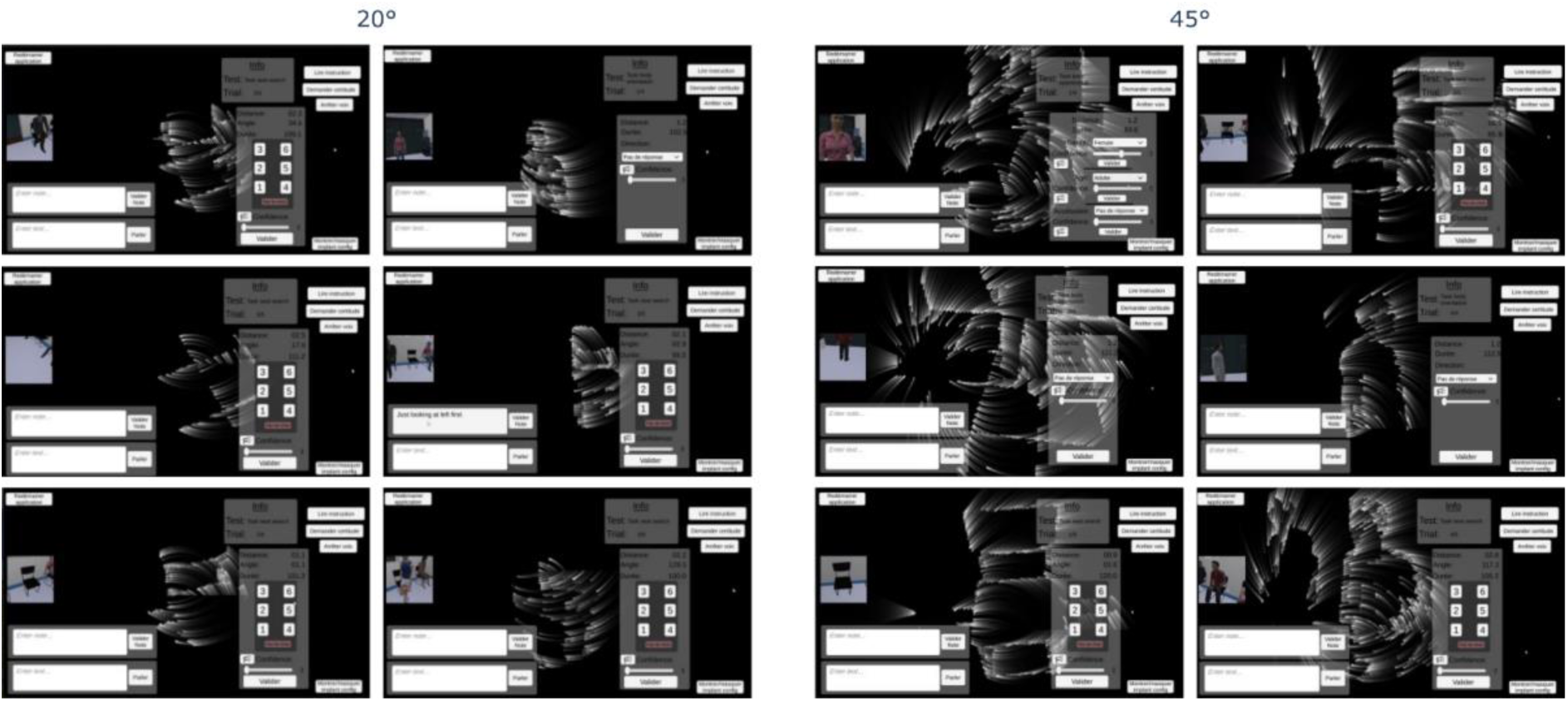
Experimenter view comparing SAV 20° (left) and SAV 45° (right) viewing conditions.

### 2.3 VR Space

The experimental area measured 4.15 m × 6.30 m (**Figure 2a**). The virtual environment simulated a medical practice, offering a realistic setting for the tasks. The 3D environment was created using assets from a 3D library, with the room designed to accurately reflect the actual physical space where the experiment took place. This careful mapping maximized the available area while ensuring participant safety, and allowed for the study of natural movement without relying on teleportation or controller-based translation. Virtual human characters were sourced from the Rocketbox library [30], providing a diverse range of figures in terms of age, ethnicity, and occupation. These fully rigged humanoid characters support pre-recorded animations, utilize pathfinding algorithms, and display facial expressions using blendshapes. Ambient sounds mimicked the subtle background noise of a hospital, enhancing immersion and preventing unintended auditory cues from influencing exploration and spatial behavior. The experiment was implemented in Unity 2019.2.16f1.

**Figure 2.**
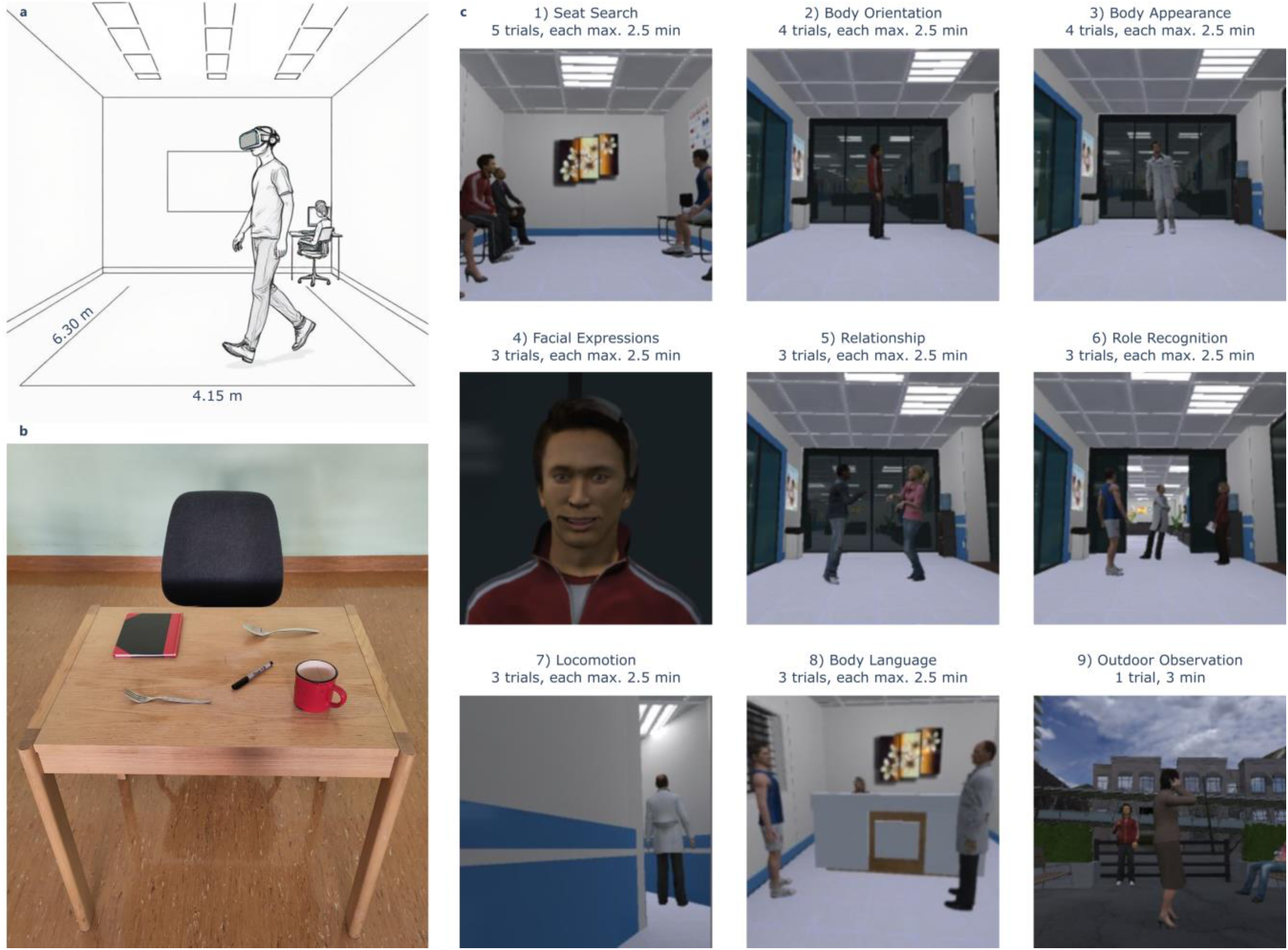
**a** Sketch of the experimental space in which participants explored a medical practice scene through a VR HMD. **b** Setup of the familiarization phase in AR. **c** Experimenter screen views of the nine tasks in the medical practice narrative.

### 2.4 Experimental protocol

Upon arrival, participants were guided to the experimental VR room. After receiving an explanation of the study and detailed instructions, participants signed the informed consent form. Participants underwent eye dominance testing using the Dolman method, put on the HMD and the standard eye tracking calibration was performed. Then, a 10-min long familiarization phase with the assigned condition (SAV 20° or SAV 45°) started. During this phase, participants completed three tasks designed to train general identification skills without directly relating to the study tasks (**Figure 2b**). The familiarization was conducted in AR, where the incoming image from the HMD camera was transformed into SAV in real-time. The tasks included identifying the borders of a table in front of them, recognizing the position and name of five objects on the table (cup, spoon, fork, pen, and notebook), and placing the cup on a coaster. These tasks were adapted from the familiarization procedure of a previous study [21]. Following familiarization, participants entered the experimental arena, where they navigated to the starting position, marked by a white space on the floor. Once correctly positioned, they were instructed to look at another white space on the opposite side of the room to calibrate both their position and gaze. Subsequently, the first set of instructions was delivered through the speakers of the HMD. All instructions are provided as **Supplementary Material**. Participants were given the opportunity to ask for clarification after instruction, ensuring full understanding before proceeding. The experiment started with a countdown once the participant was ready.

The study followed a guided social narrative consisting of a series of nine tasks, presented in the same order for all participants (**Figure 2c**). These tasks were selected based on socially relevant activities identified in previous studies [1, 31, 32], and covered a range of social skills including: localizing people in the environment, describing body orientation, analyzing physical appearance, recognizing emotions, identifying gestures, recognizing a celebrity or role, detecting motion, and analyzing body language. Participants were instructed to imagine that they were attending a medical appointment. The medical practice setting was chosen as an appropriate context, as it involves well-known social protocols and roles (e.g., doctor and secretary). The tasks were presented in the following sequence. In the first task (Seat Search), participants found themselves entering the waiting room of a medical practice. They were instructed to locate the only empty chair in the room, navigate towards it, and, once they believed they had identified it, stand in front of it, point towards it, and verbally indicate their choice to the experimenter. Following the Seat Search, participants were informed that there was some waiting time and were asked to observe the people entering the room. In the second task (Body Orientation) participants focused on the body orientation of the incoming avatars and reported whether the avatar was oriented towards them, away from them, to the left, or to the right. In the third task (Body Appearance) participants were asked to verbally report the characteristics of an incoming avatar. This included identifying the avatar’s gender (male or female), age group (adult or child), and accessories (scarf, stethoscope, beanie or handbag). The fourth task (Facial Expressions) required participants to identify the facial expression of an incoming avatar as either positive, neutral, or negative. For the fifth task (Relationship) participants were asked to determine the relationship between two incoming avatars, categorizing it as either strangers, professional, or close. In the sixth task (Role Recognition) three avatars entered the waiting room, and participants were tasked with identifying the doctor. They were instructed to walk towards the avatar they identified as the doctor and point them out. In the seventh task (Locomotion) participants were required to follow the doctor to another room. The eighth task (Body Language) took place in the medical room where participants were confronted with the doctor, secretary, and another patient. They were asked to verbally report which individual the doctor was addressing (themselves, the secretary or another patient). Notably, ambient hospital background sounds provided no auditory cues regarding the positions of the individuals, meaning that participants had to rely entirely on their visual perception. Finally, in the last task (Outdoor Observation), participants found themselves outside the hospital after the medical appointment had concluded. They were instructed to identify as many people and attributes in the outdoor scene as possible, including posture (sitting, standing, or lying), age group (adult or child), and gender (male or female). Participants were allowed to move and approach objects or avatars as they felt necessary. For some tasks (Seat Search, Body Orientation, Body Appearance, Facial Expressions, Relationship, Role Recognition, and Body Language), participants were asked to indicate their confidence level on a scale from 0 (not at all confident) to 5 (very confident) aloud once a choice was made. Participants were instructed not to provide an answer if they were unsure and were simply guessing.

Each task was repeated a varying number of times (indicated in **Figure 2c**). The number of trials for each task was determined based on a balance between several factors. Specifically, for every task, each realization (e.g., positive, negative or neutral for Facial Expression) should be presented the same number of times, but at the same time, the total number of trials needed to be kept within a reasonable range to prevent the experiment from becoming overly long. Pilot testing revealed a number of trials for each task that provided a good balance between ensuring adequate data collection and preventing participant fatigue. To ensure that participants did not rely on previous trial realizations to guess the answer in subsequent trials (e.g., identifying neutral emotion after having seen positive and negative emotions in previous trials), participants were instructed that the realizations were completely randomized and could be repeated and were explicitly told not to use any information from previous trials when solving the current one. After the study, participants confirmed that they followed these instructions. The order of task realizations was randomly shuffled between repetitions. For the first eight tasks, a maximum trial time of 2.5 minutes was set, and the final task (Outdoor Observation) had a 3-minute duration. These times were imposed to prevent task saturation, maintain comparability between tasks, and ensure participants completed tasks within reasonable time frames as the goal was to assess whether tasks could be completed in practical, natural time scales. Pilot testing showed that if participants couldn’t finish within 2.5 minutes, they likely wouldn’t have completed it in more time, making this limit appropriate. While participants could end the first eight tasks early upon making a choice, the Outdoor Observation task always lasted the full 3 minutes, as participants were tasked with identifying as many features as possible. Between every trial and task, the HMD would return to the VR calibration room, where participants were required to walk back to the starting position for the upcoming trial to recalibrate. During these inter-trial periods, participants were allowed to request breaks, and at least one break was always given in the middle of the experiment. Before the first trial of each task, the instructions for the new task were played, and a countdown timer for the new trial began. For all tasks, participants were instructed to complete tasks as quickly and accurately as possible. Throughout the experiment, the experimenter remained present in the VR room to enter results, monitor the study, and ensure participant safety. The experimenter used a control software built in Unity (version 2019.2.16f1) to record participants’ choices, confidence ratings, and timestamps of decision times.

After the study, a debriefing was conducted allowing participants to ask questions about the study.

### 2.5 Data Collection and statistical analysis

Participant head position and rotation were recorded using the HMD at 50 Hz. The positions of VR avatars and additional objects of interest were also recorded. For each task and trial, a CSV file was generated containing frame-by-frame position, rotation, and gaze data for both the participant and relevant task objects. Additionally, a separate CSV file was created for each trial and task, documenting task performance data entered by the experimenter. This file included the participant’s choice or performance outcome, as well as decision time and reported confidence for indoor identification tasks. The experimenter was not blinded to the trial conditions.

Data preprocessing was conducted in Python (version 3.9.1), and all statistical analyses were performed in R (version 4.1.1). The statistical procedures varied based on the task category and the measured variables. For the indoor identification tasks, four variables were analyzed: choice rate, success rate, decision time and confidence (the two last ones both for all trials and for successful trials only). Generalized linear mixed-effects models (GLMMs) were used to analyze choice rate, success rate, and decision time across tasks and viewing conditions. These models included viewing condition, task, and their interaction as fixed effects, and participants as random effects, with an additional offset term to adjust for the varying number of trials between tasks. Model selection followed a structured procedure: standard GLMMs (including respective fixed and random effects, trial offset, and appropriate distribution family) and models that additionally accounted for zero-inflation and overdispersion were specified using the *glmmTMB* package [33]. The models were compared using differences in Akaike’s Information Criterion (dAIC) via the *AICtab* function. The model with the lowest dAIC was tested for violations of assumptions, including correct distribution, dispersion, and homogeneity of variance of residuals, assessed using the *simulateResiduals* function from the *DHARMa* package [34]. If assumptions were not met, the model with the next lowest dAIC was tested iteratively until a suitable model was identified. If no model met all assumptions, the model with the fewest violations was selected, as linear mixed models are robust to distributional assumption violations [35]. For confidence ratings, a cumulative link mixed model (CLMM) with a logit link function was applied using the *ordinal* package, given the ordinal nature of the variable. The model specification for fixed and random effects followed the same approach as for the other measures. For significant fixed effects of viewing condition and task, Holm-adjusted post hoc comparisons were conducted using the *emmeans* package to identify specific task-related differences in viewing conditions. Statistical significance was always set to p < 0.05. The Locomotion task measured success rate, time spent outside the corridor (time in wall), mean distance to the doctor, and smoothness of this distance (operationalized as the standard deviation of this distance over the course of the trial). Generalized linear mixed-effects models were fitted with a binomial distribution for success and a Gaussian distribution for the remaining variables. Viewing condition was included as a fixed effect, and participants as a random intercept to account for repeated measures. The same model selection and assumption-checking procedures as described for the indoor identification tasks were applied. Since the Outdoor Observation task consisted of a single trial per participant, no repeated measures tests were required. Homogeneity of variance was assessed using Levene’s test, and normality was tested using the Shapiro-Wilk test. If assumptions were met, group differences between SAV 20° and SAV 45° viewing conditions were analyzed using a two-sided independent samples t-test. Otherwise, a Mann-Whitney U test was performed. In all tasks, for significant differences between the SAV 20° and SAV 45° conditions, effect sizes were computed as follows: Cohen’s d for normally distributed variables (using the *effectsize* package), rate ratios for binary outcome variables, count data, or positive continuous variables best described by binomial or Tweedie distributions, and odds ratios for ordinal logistic regression models assessing confidence. All models and assumption tests are detailed in the **Supplementary Material**.

## 3. RESULTS

To assess the impact of visual angle (SAV 20° vs. SAV 45°) on social interaction, we categorized the nine tasks into three categories. First, we looked at indoor identification tasks (Seat Search, Body Orientation, Body Appearance, Facial Expressions, Relationship, Role Recognition, and Body Language), which involved recognizing socially relevant aspects in the medical scene. Then, we looked at the Locomotion task, focusing on the ability to follow a person accurately and safely. Last, we analyzed the Outdoor Observation task, assessing the ability to localize and identify individuals and their characteristics in a crowded outdoor environment. For indoor identification tasks, we evaluated decision-making performance (choice rate, decision time, and confidence) and choice-accuracy performance (success rate, decision time in successful trials, and confidence in successful trials). For Locomotion, we assessed the ability to follow a person based on success rate, time spent outside the designated corridor (i.e., in the wall), and both the mean and smoothness (standard deviation) of their trajectory distance to the doctor. Finally, for the Outdoor Observation task, we examined the number of identified individuals and an attribute score summarizing correctly named features of these individuals.

### 3.1 A Wide visual angle facilitates decision-making in indoor identification tasks

Decision-making is defined as the ability to reach a conclusion before the trial time expires. To assess decision-making performance in indoor identification tasks, we computed three key measures. Choice rate quantifies the number of trials in which a participant reached a conclusion (**Figure 3a**). Decision time is the mean time taken to make a decision in trials where a conclusion was reached (**Figure 3b**). Confidence reflects the mean confidence rating for trials where a choice was made (**Figure 3c**).

**Figure 3.**
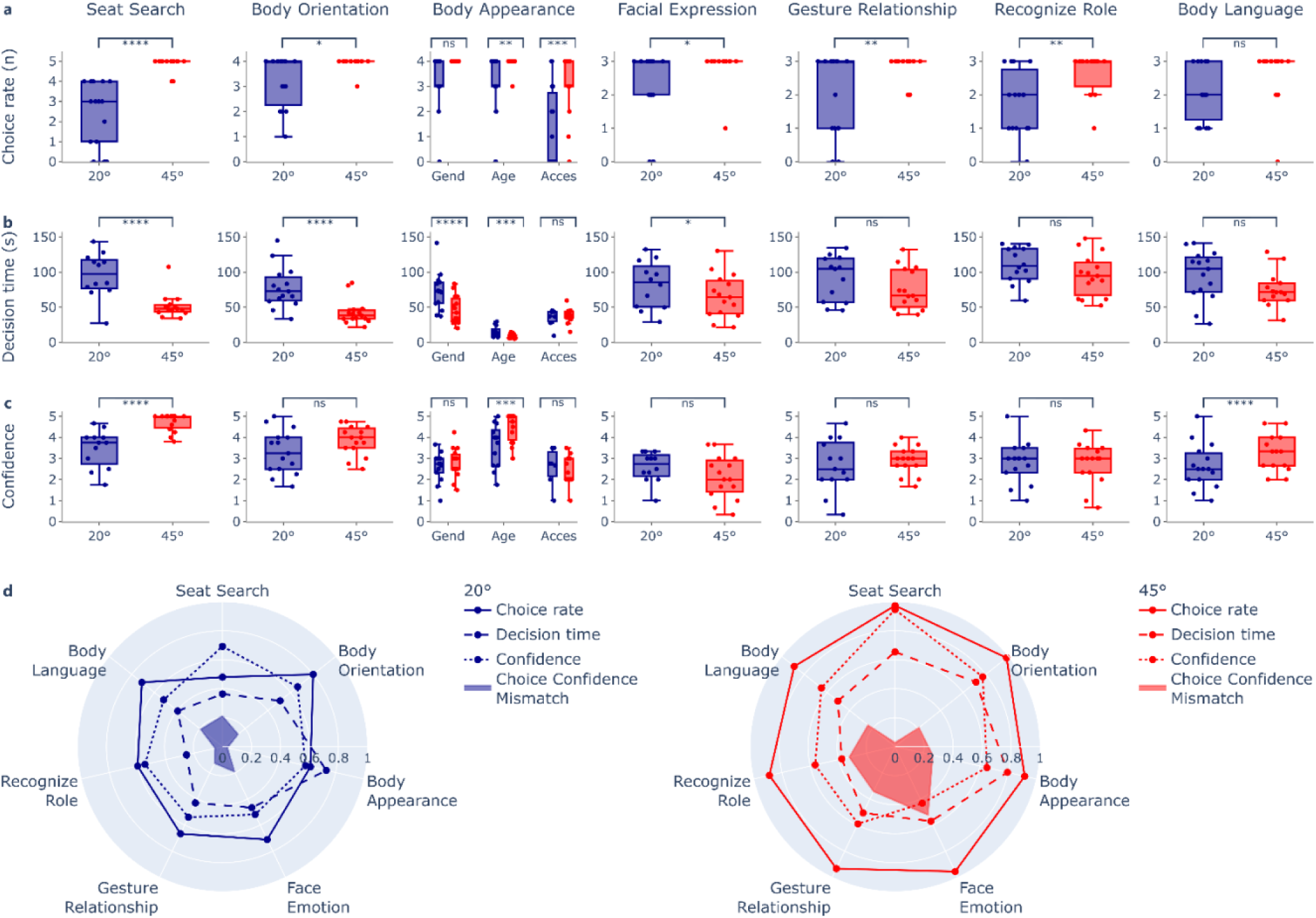
Decision-making performance as a function of the SAV viewing condition for all indoor identification tasks. (a) Quantification of choice rate, (b) decision time and (c) confidence ratings for task performance for all indoor identification tasks. Boxes span the interquartile range (IQR) from the 25^th^ percentile to the 75^th^ percentile, the line inside represents the median, and the whiskers extend to the most extreme data points within 1.5 times the IQR. Outliers are data points falling beyond 1.5 times the IQR. P-values indicate two-tailed post hoc z-tests on the respective generalized linear mixed effects models and are reported as: ns p > 0.05, * p < 0.05, ** p < 0.01, *** p < 0.001, and **** p < 0.0001. (d) Summary plot for decision-making performance at 20° (left) and 45° (right), displaying normalized metrics for choice rate (solid line), decision time (dashed line), confidence (dotted line), and choice-confidence mismatch (shaded area).

A significant main effect of viewing condition and task was found for choice rate (viewing condition: *χ*² = 20.60, p < 0.0001, df = 1; task: *χ*² = 85.56, p < 0.0001, df = 8; two-tailed ANOVA on generalized linear mixed effects model with Binomial distribution), allowing further investigation of viewing condition differences within tasks. Participants in the SAV 45° condition reached significantly more conclusions than those in SAV 20° across nearly all tasks, except for Body Appearance Gender and Body Language (**Figure 3a**; Seat Search: z = -4.73, p < 0.0001, RR = 0.01; Body Orientation: z = -2.41, p = 0.0159, RR = 0.05; Body Appearance Gender: z = -0.004, p = 0.9966; Body Appearance Age: z = -2.75, p = 0.0059, RR = 0.04; Body Appearance Accessory: z = -3.87, p = 0.0001, RR = 0.07; Facial Expressions: z = -2.54, p = 0.0110, RR = 0.08; Relationship: z = -2.62, p = 0.0088, RR = 0.10; Role Recognition: z = -2.63, p = 0.0086, RR = 0.12; Body Language: z = -1.73, p = 0.0837; two-tailed post hoc z-tests).

For trials in which a choice was made, a significant main effect of viewing condition and task was found for decision time (viewing condition: *χ*² = 25.08, p < 0.0001, df = 1; task: *χ*² = 1485.23, p < 0.0001, df = 8; two-tailed ANOVA on generalized linear mixed effects model with Tweedie distribution). A wider visual angle SAV 45° resulted in significantly faster decisions in the Seat Search, Body Orientation, Body Appearance Gender, Body Appearance Age, and Facial Expressions tasks, but not in the others (**Figure 3b**; Seat Search: z = 5.93, p < 0.0001, RR = 1.96; Body Orientation: z = 5.18, p < 0.0001, RR = 1.80; Body Appearance Gender: z = 4.32, p < 0.0001, RR = 1.80; Body Appearance Age: z = 3.85, p = 0.0001, RR = 1.71; Body Appearance Accessory: z = -0.24, p = 0.8102; Facial Expressions: z = 2.29, p = 0.0218, RR = 1.33; Relationship: z = 1.70, p = 0.0899; Role Recognition: z = 1.72, p = 0.0851; Body Language: z = 1.83, p = 0.0680; two-tailed post hoc z-tests).

For these trials, an ordinal regression model further revealed that those in the SAV 45° viewing condition generally reported significantly higher confidence levels than participants in the SAV 20° condition (β = 3.14, SE = 0.62, z = 5.07, p < 0.0001). Post hoc pairwise comparisons indicated that this increase in confidence was, however, only significant in the Seat Search, Body Appearance Age, and Body Language tasks (**Figure 3c**; Seat Search: z = -5.07, p < 0.0001, OR = 0.04; Body Orientation: z = -1.88, p = 0.0596; Body Appearance Gender: z = -1.03, p = 0.3016; Body Appearance Age: z = -3.75, p = 0.0002, OR = 0.11; Body Appearance Accessory: z = 1.04, p = 0.2968; Facial Expressions: z = 1.59, p = 0.1127; Relationship: z = -0.52, p = 0.6037; Role Recognition: z = -0.22, p = 0.8297; Body Language: z = -0.05, p = 0.0402, OR = 0.29; two-tailed post hoc z-tests).

The summary plot (**Figure 3d**) illustrates the impact of visual angle on decision-making across tasks. Qualitatively, participants in the SAV 45° condition were able to reach a conclusion in nearly all tasks (red solid line near the maximum), whereas those in SAV 20° struggled more (blue solid line closer to 0.5). This pattern was also observed for decision time, with participants in SAV 45° reaching conclusions faster in most tasks (dashed lines). However, reported confidence levels did not consistently reflect this improvement. Except for the seat search task, confidence levels for SAV 45° participants were not visibly higher than those for SAV 20° participants (dotted lines). As a result, a greater discrepancy between choice rate and confidence emerged for SAV 45° (shaded areas), raising the question whether this mismatch between decision-making performance and confidence levels predicts choice accuracy.

### 3.2 A Wide visual angle enhances choice accuracy in indoor identification tasks

Choice accuracy is defined as the ability to reach a successful conclusion before the end of the trial time. To assess task choice-accuracy performance in indoor identification tasks, we computed three measures. Success rate is the number of trials in which participants arrived at a correct conclusion (**Figure 4a**). Decision time in successful trials measures the mean time taken to make a correct decision (**Figure 4b**). Confidence in successful trials captures the mean confidence rating when a correct conclusion was reached (**Figure 4c**).

**Figure 4.**
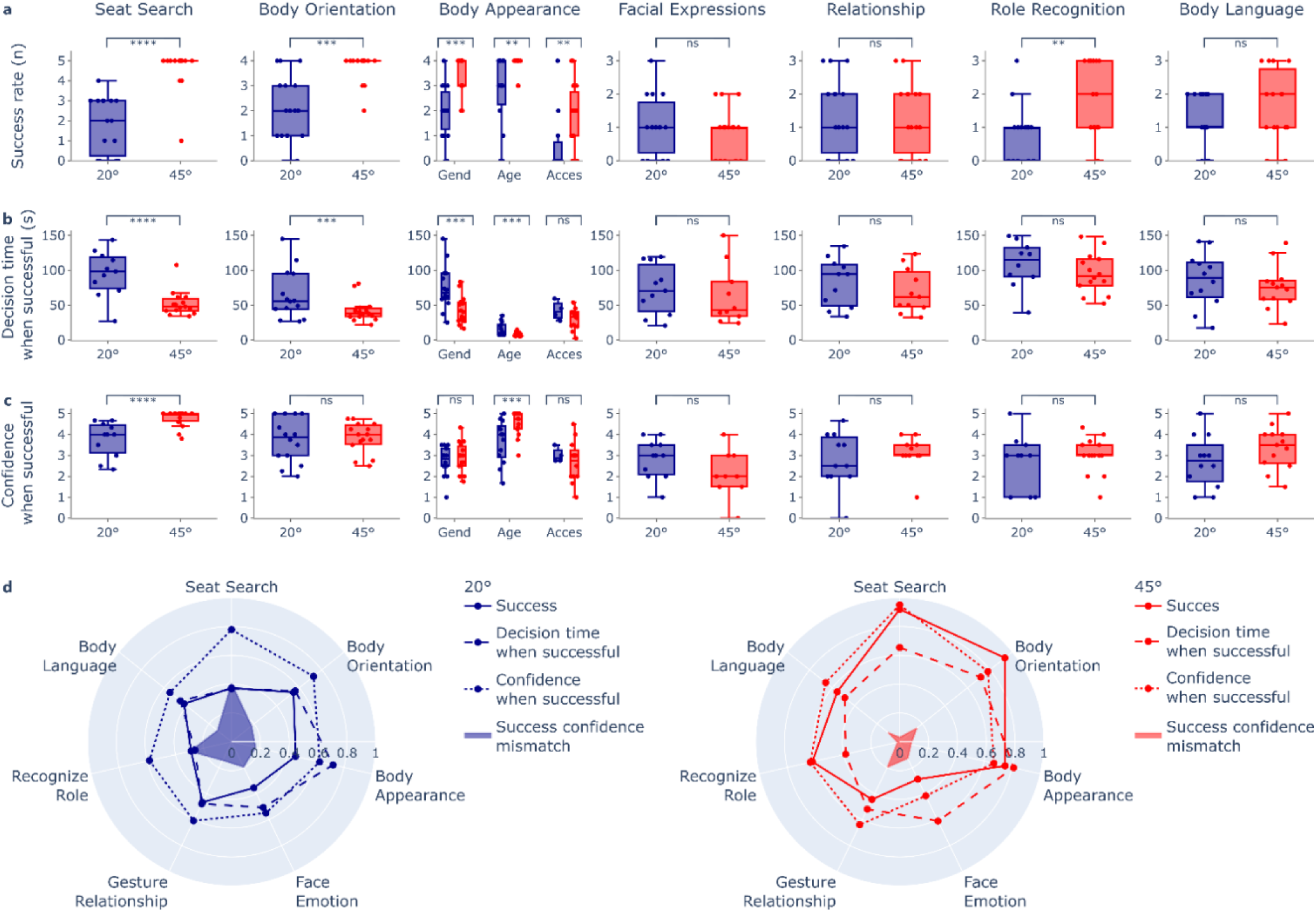
Choice-accuracy performance as a function of the SAV viewing condition for all indoor identification tasks. (a) Quantification of success rate, (b) decision time when successful, and (c) confidence ratings for successful trials across all indoor identification tasks. Boxes span the IQR from the 25^th^ percentile to the 75^th^ percentile, the line inside represents the median, and the whiskers extend to the most extreme data points within 1.5 times the IQR. Outliers are data points falling beyond 1.5 times the IQR. P-values indicate two-tailed post hoc z-tests on the respective generalized linear mixed effects models and are reported as: ns p > 0.05, * p < 0.05, ** p < 0.01, *** p < 0.001, and **** p < 0.0001. (d) Summary plot of SAV 20° (left) and SAV 45° (right), displaying normalized metrics for success rate (solid line), decision time (dashed line), confidence (dotted line), and success-confidence mismatch (shaded area).

A significant main effect of both ‘viewing condition’ and ‘task’ was observed for success rate (‘viewing condition’: *χ*² = 18.69, p < 0.0001, df = 1; ‘task’: *χ*² = 79.60, p < 0.0001, df = 8; two-tailed ANOVA on a generalized linear mixed effects model with a Binomial distribution). Further analysis of viewing condition differences within tasks revealed that a wide visual angle (SAV 45°) resulted in significantly more correct trials compared to SAV 20° in most tasks, except for Facial Expressions, Relationship, and Body Language (**Figure 4a**; Seat Search: z = -5.65, p < 0.0001, RR = 0.04; Body Orientation: z = -3.96, p = 0.0001, RR = 0.08; Body Appearance Gender: z = -3.34, p = 0.0008, RR = 0.18; Body Appearance Age: z = -3.10, p = 0.0020, RR = 0.11; Body Appearance Accessory: z = -3.03, p = 0.0024, RR = 0.22; Facial Expressions: z = 0.51, p = 0.6131; Relationship: z = 0.14, p = 0.8876; Recognize Role: z = -2.93, p = 0.0033, RR = 0.21; Body Language: z = -1.19, p = 0.2329; two-tailed post hoc z-tests).

For trials in which a correct choice was made, a significant main effect of both ‘viewing condition’ and ‘task’ was also observed for decision time (‘viewing condition’: *χ*² = 23.18, p < 0.0001, df = 1; ‘task’: *χ*² = 1225.12, p < 0.0001, df = 8; two-tailed ANOVA on a generalized linear mixed effects model with a Tweedie distribution). A wide visual angle (SAV 45°) led to significantly faster correct decisions than SAV 20° in Seat Search, Body Orientation, and Body Appearance (Gender and Age) tasks but not in other tasks (**Figure 4b**; Seat Search: z = 5.03, p < 0.0001, RR = 1.93; Body Orientation: z = 3.62, p = 0.0003, RR = 1.60; Body Appearance Gender: z = 3.89, p = 0.0001, RR = 1.68; Body Appearance Age: z = 4.05, p = 0.0001, RR = 1.74; Body Appearance Accessory: z = 1.77, p = 0.0769; Facial Expressions: z = 1.60, p = 0.1093; Relationship: z = 0.96, p = 0.3359; Recognize Role: z = 1.25, p = 0.2105; Body Language: z = 0.79, p = 0.4303; two-tailed post hoc z-tests).

For these successful trials, the ordinal regression model indicated that those in the SAV 45° condition reported overall significantly higher confidence levels than SAV 20° participants (β = 2.85, SE = 0.68, z = 4.17, p < 0.0001). However, post hoc pairwise comparisons showed that increased confidence in the SAV 45° condition was significant only in the Seat Search and Body Appearance Age tasks, with no differences in confidence observed for other tasks (**Figure 4c**; Seat Search: z = -4.17, p < 0.0001, OR = 0.06; Body Orientation: z = -0.78, p = 0.4365; Body Appearance Gender: z = -0.86, p = 0.3892; Body Appearance Age: z = -3.92, p = 0.0001, OR = 0.08; Body Appearance Accessory: z = 0.54, p = 0.5884; Facial Expressions: z = 1.79, p = 0.0738; Relationship: z = -0.81, p = 0.4155; Recognize Role: z = -1.09, p = 0.2751; Body Language: z = -1.75, p = 0.0800, two-tailed post hoc z-tests).

The summary plot (**Figure 4d**) illustrates how viewing condition influences successful completion of the trials across the indoor identification tasks. Participants in the SAV 45° condition generally achieved higher success rates (solid lines) and completed successful trials slightly faster (dashed lines) than those in the SAV 20° condition. However, overall the confidence levels remained similar between groups (dotted lines). Notably, while the mismatch between decision-making performance and confidence is greater for SAV 45° than for SAV 20° (shaded area, **Figure 3d**), the inverse pattern is observed for choice-accuracy performance (shaded area, **Figure 4d**). In the SAV 45° condition, success rates closely align with confidence levels, whereas in the SAV 20° condition, participants exhibit overconfidence relative to their actual performance. This suggests that while both groups report similar confidence, SAV 20° participants tend to overestimate their success, whereas SAV 45° participants demonstrate performance levels that match their confidence ratings.

### 3.3 A wide visual angle improves the ability to follow people

Successfully identifying individuals and their characteristics, behaviors, and positions is essential for daily social interactions. However, beyond mere identification, another critical skill is the ability to smoothly track moving individuals and follow their trajectories. To assess this ability, we evaluated the Locomotion task, designed to measure participants’ capacity to track and follow the doctor’s position over time as they moved through a corridor (**Figure 5a,b**). Four measures were computed in this task. First, the success rate quantified the number of trials in which participants correctly followed the doctor’s trajectory (**Figure 5c**). Second, we analyzed participants’ spatial accuracy by examining the time spent outside the designated corridor, defined as ‘time in wall’, which reflects how much participants deviated from the correct path (**Figure 5d**). Third, we assessed participants’ ability to maintain proximity to the doctor over the course of the trial (**Figure 5e**) by computing their average distance to the doctor throughout the trial (**Figure 5f**). Lastly, we quantified the variability in this distance over time using the standard deviation of the doctor-participant distance as a measure of tracking smoothness (**Figure 5g**).

**Figure 5.**
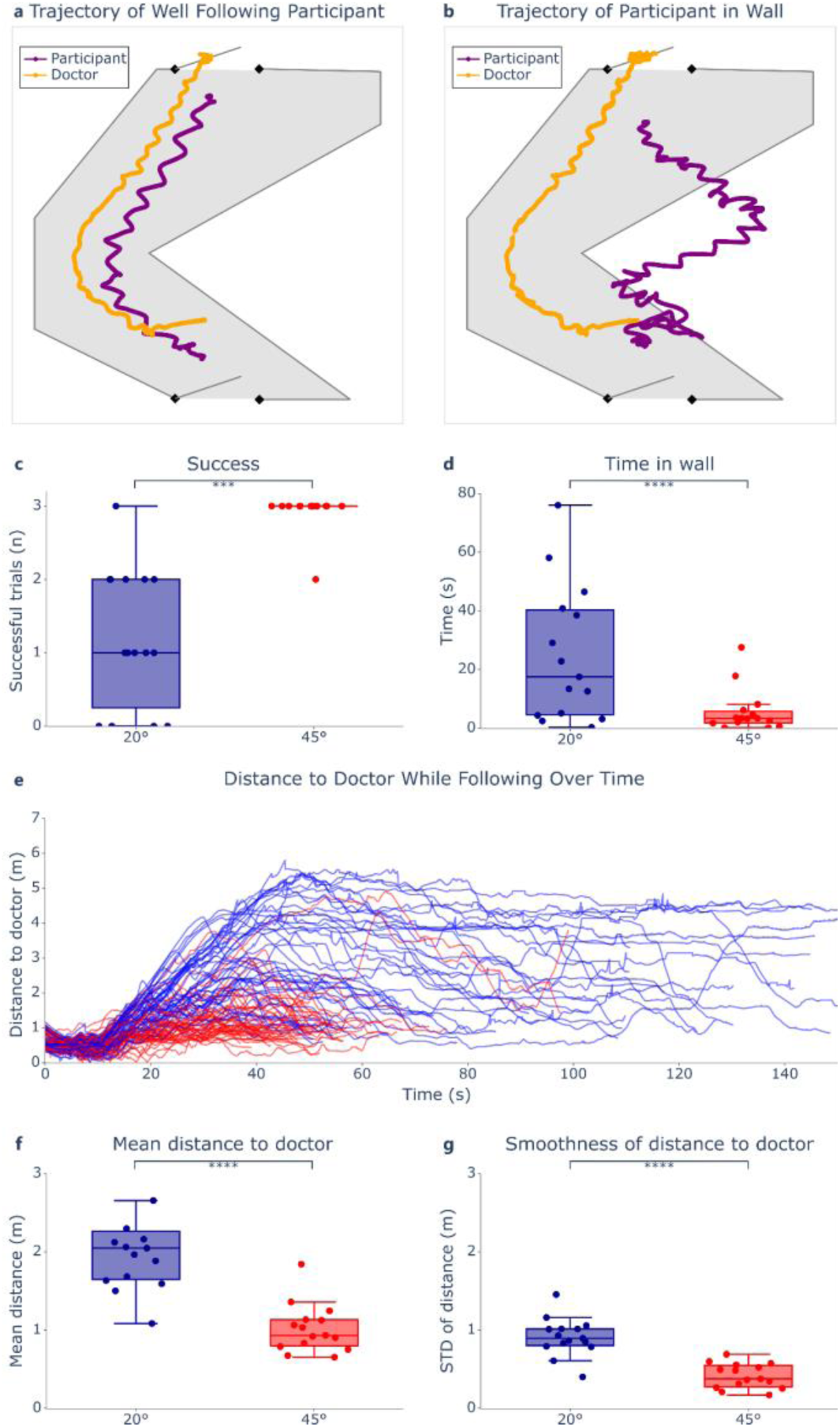
Locomotion task performance as a function of the SAV viewing condition. (a) Schematic of a participant (purple line) who followed the doctor (yellow line) well. (b) Example of a participant’s trajectory deviating from the corridor. Quantification of (c) success rate and (d) time spent outside the corridor. (e) Participants’s distances to the doctor over time. Every line represents one trial (blue: SAV 20°, red: SAV 45° participants). (f) Average distance to the doctor. (g) Distance smoothness quantified as standard deviation of the doctor-participant distance. Boxes span the IQR from the 25 ^th^ percentile to the 75^th^ percentile, the line inside represents the median, and the whiskers extend to the most extreme data points within 1.5 times the IQR. Outliers are data points falling beyond 1.5 times the IQR. P-values indicate the viewing condition fixed effects of the respective generalized linear mixed effects models and are reported as: ns p > 0.05, * p < 0.05, ** p < 0.01, *** p < 0.001, and **** p < 0.0001.

A generalized linear mixed-effects model with a Binomial distribution revealed that participants in the SAV 45° condition were significantly more likely to successfully follow the doctor compared to those in the SAV 20° condition (**Figure 5c**: β = -4.33, SE = 1.15, z = -3.77, p = 0.0002, RR = 0.0131). In addition, participants in the SAV 20° condition spent significantly more time outside the corridor, indicating greater difficulty in maintaining an accurate trajectory (**Figure 5d**: β = -46.53, SE = 10.04, z = -4.64, p < 0.0001, Cohen’s d = 0.65). Furthermore, participants in the SAV 45° condition were able to follow the doctor more closely (**Figure 5f**: β = -1.08, SE = 0.16, z = -6.72, p < 0.0001, Cohen’s d = 1.53) and with greater smoothness, demonstrating less variability in their distance to the doctor over time (**Figure 5g**: β = -0.50, SE = 0.07, z = -6.79, p < 0.0001, Cohen’s d = 1.41). These findings suggest that a wider visual angle facilitates not only the ability to detect and track moving individuals but also enables smoother and more precise following behavior.

### 3.4 A cluttered outdoor environment limits the advantage of a wide visual angle on person recognition

Lastly, we examined whether the difference between SAV 20° and SAV 45° viewing conditions persisted in a cluttered outdoor environment. To assess this, we quantified two measures in the Outdoor Observation task: (1) the number of identified peopleand (2) the attribute score, calculated as the total number of correctly identified attributes across all avatars.

Qualitatively, participants in the SAV 45° condition tended to identify more people and correctly report more attributes than those in the SAV 20° condition (**Figure 6**). However, statistical comparisons did not reach significance. A Shapiro-Wilk normality test revealed that the number of identified people in the SAV 20° condition significantly deviated from normality (Shapiro-Wilk SAV 20°: W = 0.87, p = 0.0321; Shapiro-Wilk SAV 45°: W = 0.90, p = 0.0815; Levene’s test: F = 0.01, p = 0.9091). Consequently, we conducted a Mann-Whitney U test, which showed no significant difference between the two viewing conditions in terms of the number of people identified (**Figure 6a**: W = 86, p = 0.2585). For the attribute score, assumptions of normality and homogeneity of variance were met (Shapiro-Wilk SAV 20°: W = 0.8966, p = 0.0846; Shapiro-Wilk SAV 45°: W = 0.9177, p = 0.1775; Levene’s test: F = 0.01, p = 0.9091). A two-sided independent samples t-test revealed a statistically non-significant increase in attribute identification for the SAV 45° condition compared to SAV 20° (**Figure 6b**: t(28) = -1.60, p = 0.1217). These results suggest that in a cluttered outdoor environment, the number of identified people and the attribute score did not significantly differ between the SAV 20° and SAV 45° viewing conditions.

**Figure 6.**
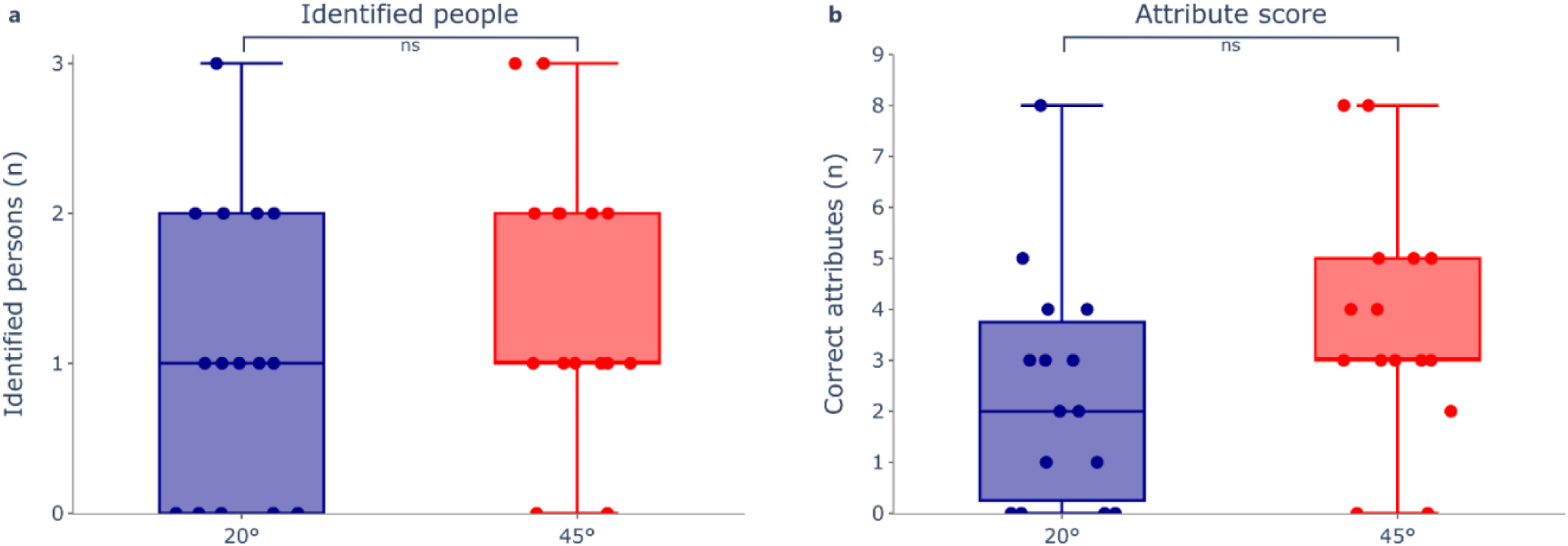
Person recognition performance in a cluttered outdoor environment as a function of the SAV viewing condition. (a) Quantification of number of identified people and (b) attribute score. Boxes span the IQR from the 25^th^ percentile to the 75^th^ percentile, the line inside represents the median, and the whiskers extend to the most extreme data points within 1.5 times the IQR. P-values indicate results from the Mann-Whitney U (a) and the two-sided independent samples t-test (b) and are reported as: ns p > 0.05, * p < 0.05, ** p < 0.01, *** p < 0.001, and **** p < 0.0001.

## DISCUSSION

Social interaction is largely visual, with nonverbal cues essential for communication and social integration. Face recognition and social perception remains a major challenge for visually impaired individuals, impacting relationships and engagement [5]. While compensatory strategies help, they cannot fully replace visual processing, and no assistive technology yet restores real-time social perception, underscoring the need to optimize artificial vision [10]. Despite the critical role of vision in social interaction, research on artificial vision has largely focused on object recognition, spatial navigation, and mobility, with social perception remaining an underexplored domain. Existing studies on low vision and blindness have documented impairments in recognizing faces, emotions, and body language but are often limited to questionnaire-based assessments or controlled tasks without realistic social dynamics [1, 36]. In prosthetic vision, research on social perception has focused on static face recognition and small-scale controlled experiments [37, 38]. SAV studies, while offering controlled manipulation of perceptual parameters, have been largely confined to navigation and object perception tasks. This study addresses these gaps by investigating how visual field size influences social perception in a VR-based medical practice scenario. Unlike prior research, it evaluates a broad range of social tasks within a unified framework, enabling comparisons across different aspects of social interaction. Moreover, it integrates immersive storytelling and realistic SAV modeling grounded in real implant parameters, phenomenological observations, and anatomical data [16, 25, 27, 39]. By extending artificial vision research beyond object recognition and mobility, this study provides critical insights into the functional relevance of visual field expansion for social interactions in the real-world.

Participants in the SAV 45° condition exhibited significantly higher choice rates and faster decision-making across all indoor identification tasks, reinforcing the role of an expanded visual field in facilitating efficient visual exploration. These findings align with prior research showing that a minimum of 20° to 35° is required for natural task solving and that a broader field enhances object detection, reduces scanning effort, and accelerates processing time [15, 40]. The present study extends this research by demonstrating that an expanded field improves not only spatial awareness but also social perception in structured indoor settings. Beyond improving decision speed, the SAV 45° condition also enhanced task accuracy, aligning with evidence that increasing the number of simultaneously perceivable elements boosts performance, provided perceptual capacity is not exceeded [41]. The expanded visual angle particularly facilitated better spatial integration of social cues, improving the recognition of individuals’ locations and body orientation. Additionally, perception of attributes such as gender, age group, accessories, and social roles was enhanced. However, fewer performance gains were found for tasks requiring finer feature discrimination, such as identifying facial expressions, relationship types, or subtle body language cues, such as determining whom someone is looking at. While the SAV 45° condition substantially improved decision-making speed and accuracy compared to the SAV 20° condition, performance remained below natural vision levels. Although choice-making frequency neared its maximum across indoor tasks (**Figure 3d**), accuracy did not reach the near-perfect levels expected in sighted individuals (**Figure 4d**). This suggests that while expanding the visual field enhances spatial awareness and efficiency, it does not fully compensate for remaining limitations of artificial vision. Factors such as the elongated shape of phosphenes due to axonal activation, slow temporal processing, and perceptual fading likely contribute to this gap by increasing cognitive demands and reducing scene stability [15, 16, 42–44]. Additionally, the monocular nature of SAV, reflecting real-world prosthetic vision, might have impaired depth perception [45, 46]. However, prior research indicates that adaptation and training enhance artificial vision performance. Studies on prosthesis users show that perceptual learning improves spatial awareness and task efficiency over time [39, 47], while simulated prosthetic vision studies suggest that repeated exposure further benefits users with a wide visual angle [21]. Thus, while a 45° field provides advantages that don’t quite reach natural performance yet, its long-term impact may be greater as users refine their ability to process social and spatial cues in dynamic environments. Interestingly, confidence levels did not always reflect objective improvements. Participants in the SAV 45° condition exhibited a greater discrepancy between decision-making performance and confidence than those in the SAV 20° condition. While success rates in the SAV 45° condition closely matched reported confidence, participants in the SAV 20° condition tended to overestimate their accuracy. This suggests that individuals with a restricted visual field may rely more on heuristic-based judgments, leading to overconfidence despite lower actual accuracy. These findings align with prior research indicating that visual uncertainty prompts compensatory inferential strategies, sometimes resulting in miscalibrated confidence estimates [48]. In the locomotion task, the wider visual field significantly improved participants’ ability to track and follow a moving individual while maintaining spatial accuracy. These results are consistent with studies on low vision and mobility, which report difficulties in locomotion under restricted visual fields [49–51]. Previous findings on retinal prostheses and SAV suggest that limited fields force users to rely on compensatory scanning strategies, which introduce delays and increase cognitive burden [16, 39]. The present study extends these insights by demonstrating that within a structured indoor setting, an expanded visual field enables smoother and more accurate tracking of another person’s movement, likely by reducing the need for constant reorienting.

While the advantages of a wider visual angle were evident in structured indoor settings, they were less pronounced in a cluttered outdoor visual task. Although participants in the SAV 45° condition showed a descriptive improvement in person recognition, this advantage did not reach statistical significance. A factor that likely contributed to this diminished effect is that visual clutter imposes a greater demand on figure-ground segmentation. In structured indoor environments, limited overlapping stimuli reduce perceptual competition, whereas outdoor settings introduce a high density of competing elements, making segmentation more challenging [52]. Unlike natural vision, and unless specifically added, artificial vision lacks background suppression mechanisms, leading to information overload rather than improved recognition [53]. A wider visual field increases the amount of available visual input, but if perceptual resources are already strained by artificial vision’s axon tails and perceptual fading, additional information does not necessarily enhance recognition. Although results did not reach statistical significance, the observed trend suggests a potential advantage of a wider visual field in cluttered environments. However, additional visual processing strategies, such as adaptive contrast enhancement or dynamic segmentation algorithms, may be necessary to fully realize these benefits [54, 55].

Some limitations of the present study should be noted. First, while our tasks were designed to approximate naturalistic social interactions, they represent only a subset of real-world scenarios. Real-life interactions are highly dynamic, involving continuous multisensory feedback and active engagement, which our study did not fully capture. Nevertheless, we aimed to cover a broad range of relevant skills for social perception and interaction, grounding our task selection in established literature [1, 31, 32]. Second, while task comparability within our study is inherently constrained, several measures were implemented to mitigate this issue. Variations in the number of trials across tasks were accounted for in the statistical model using trial offsetting. Additionally, participants were explicitly instructed not to guess, eliminating the possibility of a forced-choice effect. Lastly, they were instructed that task realizations (e.g., positive, negative, or neutral facial expressions) were fully randomized, ensuring that previous trials would not be used to predict subsequent answers. Third, the SAV model, while grounded in known prosthetic vision parameters, remains an approximation. SAV was based on the POLYRETINA retinal implant, chosen for its high-resolution and wide-field artificial vision and because most knowledge about artificial vision stems from retinal implants, whereas research on other implants and optogenetics remains limited [56–62]. While the simulation incorporated key perceptual properties such as phosphene shape variability, brightness fluctuations, and perceptual fading [16, 28], it did not account for individual variability in phosphene mapping or higher-order perceptual phenomena such as spatial distortions and adaptation effects [63]. Even though the simulation was based on the POLYRETINA implant, the findings extend beyond its specific parameters, as this study focuses on general visual properties essential for effective perception and interaction in social contexts. Identifying these key parameters is critical for all vision restoration therapies, and the use of SAV in naturalistic social settings provides broader insights into artificial vision utility.

Future work should incorporate real-time interactions, as the current setup focused on passive interpretation of social cues. Introducing interactive scenarios, where participants engage in dynamic conversations, respond to gestures, or navigate social exchanges, would provide a more comprehensive assessment of artificial vision’s applicability in real-world social settings. New studies which investigate how factors beyond visual field size, such as temporal modulation or adaptive contrast enhancement, contribute to functional improvements are required. Further research should explore the integration of figure-ground segmentation algorithms and attentional mechanisms to optimize information extraction in visually cluttered environments. Lastly, long-term studies are needed to examine user adaptation to artificial vision over time and whether training enhances the ability to utilize artificial vision for social interaction. Addressing these aspects will help refine artificial vision technologies to improve social functioning and mobility for visually impaired individuals.

## Data and code availability statement

The authors declare that the data supporting the findings of this study are available in the paper. Any additional requests for information can be directed to the corresponding author. The SAV code is accessible online (https://github.com/lne-lab/polyretina_vr_social).

## Ethical statement

The research was conducted in accordance with the principles embodied in the Declaration of Helsinki. The study was approved by the Commission cantonale (VD) d’éthique de la recherche sur l’être humain (CER-VD) in Switzerland (Project-ID 2023-01770). All participants gave written informed consent to participate in the study.

## Author contribution statement

S.H. designed the experiment, collected the data, analyzed the data and wrote the manuscript. Y.T. designed and programmed the experiment and discussed the results and contributed to the manuscript. E.S. designed the experiment and discussed the results. D.G. designed and led the study, discussed the results and edited the manuscript.

## Acknowledgment

The authors would like to thank Inés Roque Avilés and Théophine Gurlie for their help in designing the experiment. This project has received funding from the European Union’s Horizon 2020 research and innovation programme under the Marie Skłodowska-Curie Grant Agreement No. 861423.

## SUPPLEMENTARY TABLES

**Supplementary Table 1.** Outdoor Observation Attribution Score Evaluation Grid.

| Evaluation Criteria | Score |
| --- | --- |
| Lady on smartphone |  |
| Gender: Female | 0: not identified; 1: correctly identified |
| Age group: Adult | 0: not identified; 1: correctly identified |
| Posture: Standing | 0: not identified; 1: correctly identified |
| Smoking man |  |
| Gender: Male | 0: not identified; 1: correctly identified |
| Age group: Adult | 0: not identified; 1: correctly identified |
| Posture: Standing | 0: not identified; 1: correctly identified |
| Lady sitting on bench |  |
| Gender: Female | 0: not identified; 1: correctly identified |
| Age group: Adult | 0: not identified; 1: correctly identified |
| Posture: Seated | 0: not identified; 1: correctly identified |

## SUPPLEMENTARY NOTES

### Instructions before each task

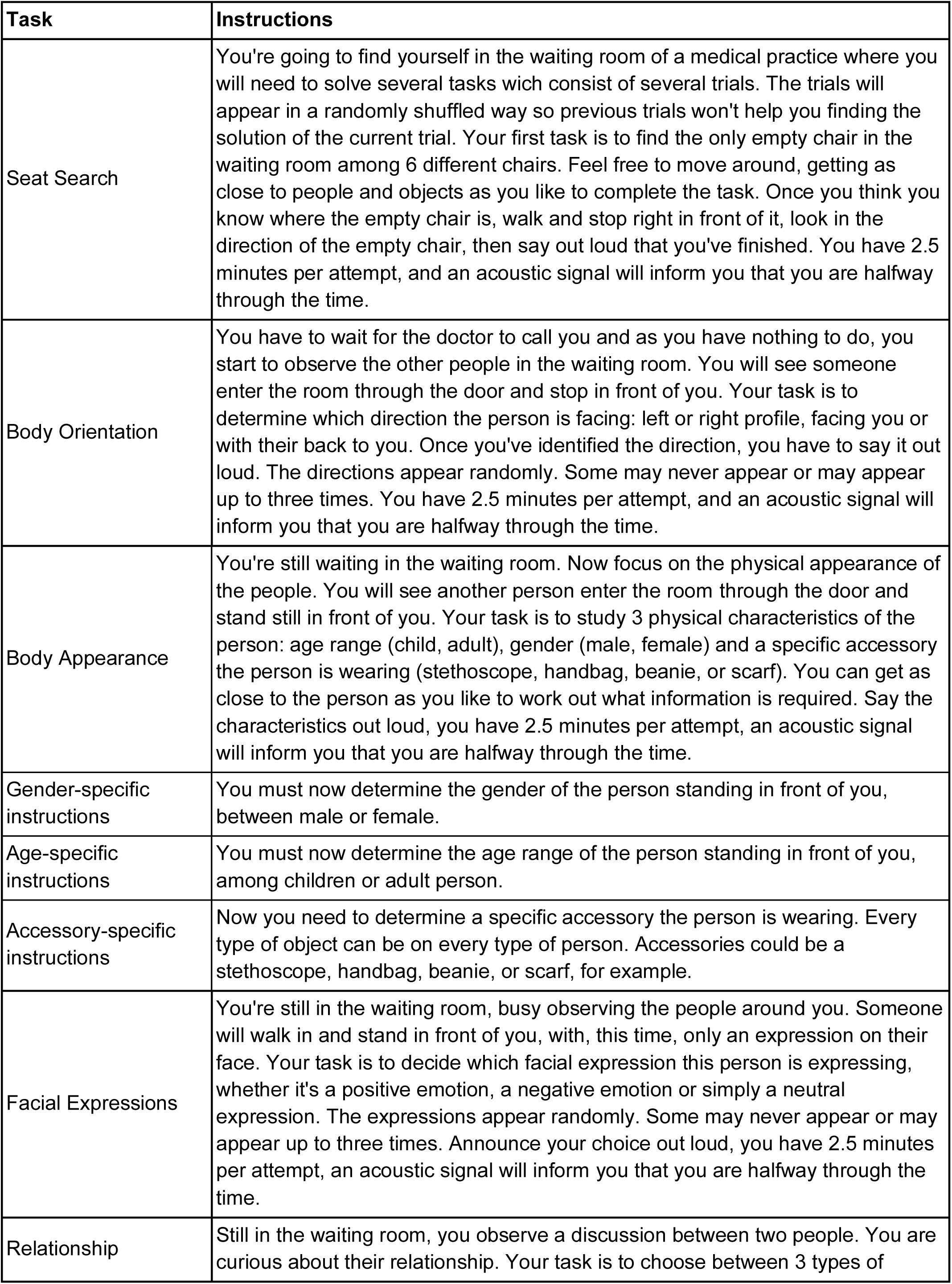
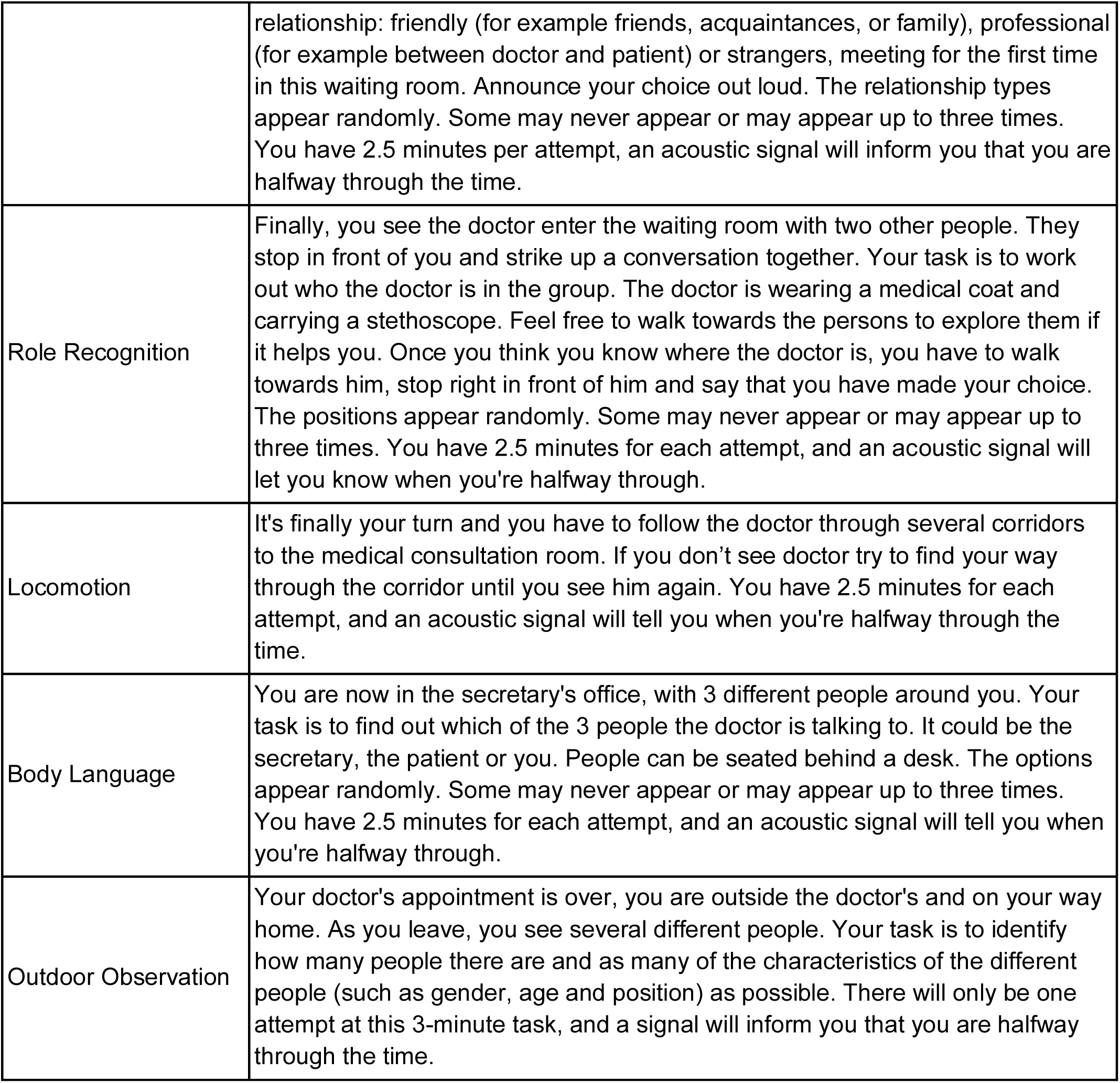

### Confidence instructions

How confident are you about your choice? (0 not sure at all, 5 very sure).

### End of experiment instructions

Thank you for taking part in this experience. You can now remove your virtual reality headset.

### Model specifics, assumptions and statistics

- Indoor identification tasks: Choice rate

Best fitting model: Generalized linear mixed-effects model with a binomial distribution. The dependent variable was the choice rate, with user condition, task, and their interaction as fixed effects. Since the design involved repeated measures, participant ID was included as a random effect. A log offset for the total number of trials was applied to account for differences in trial counts across tasks.

Model assumptions: assumption violations are highlighted in red.

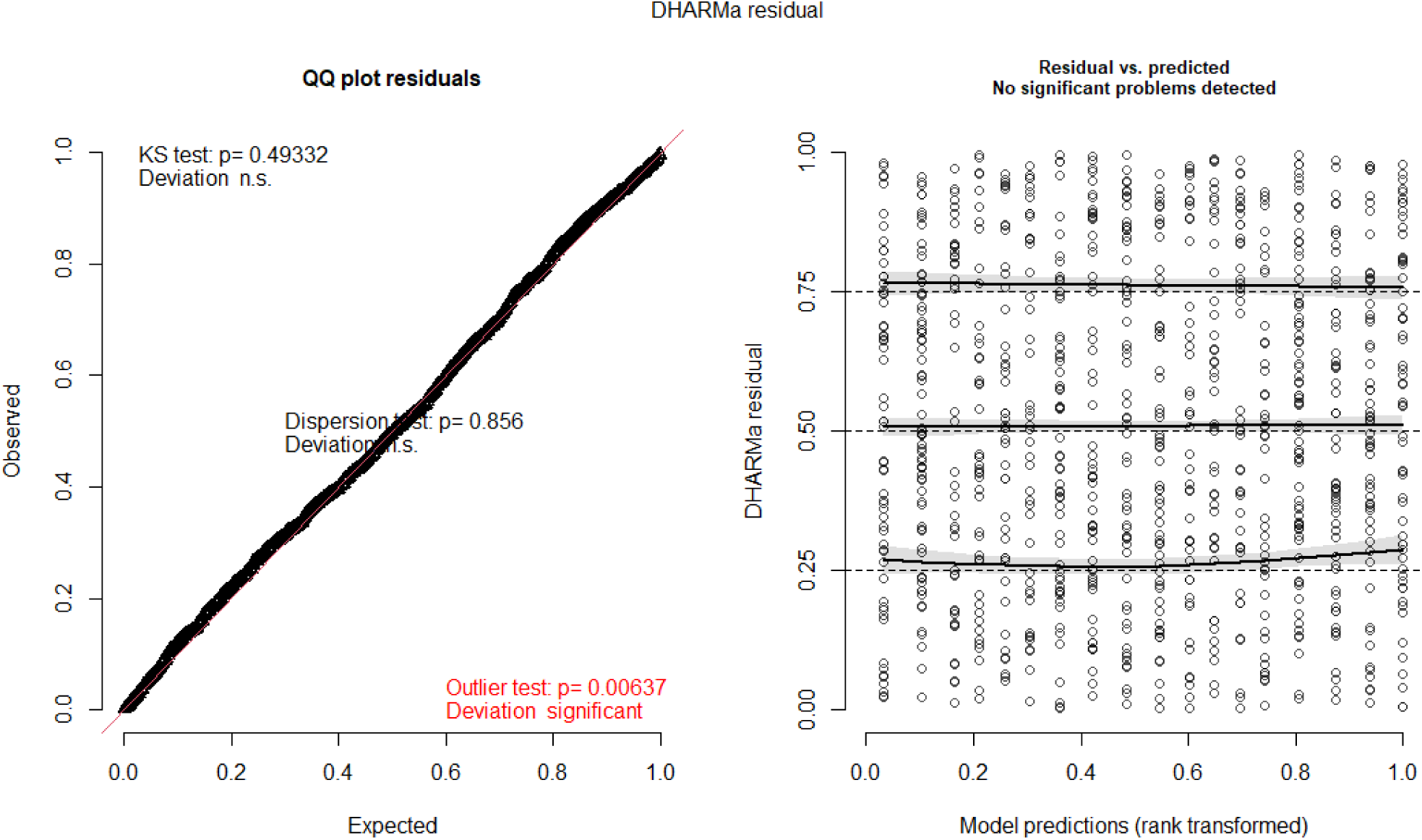

Statistical results:

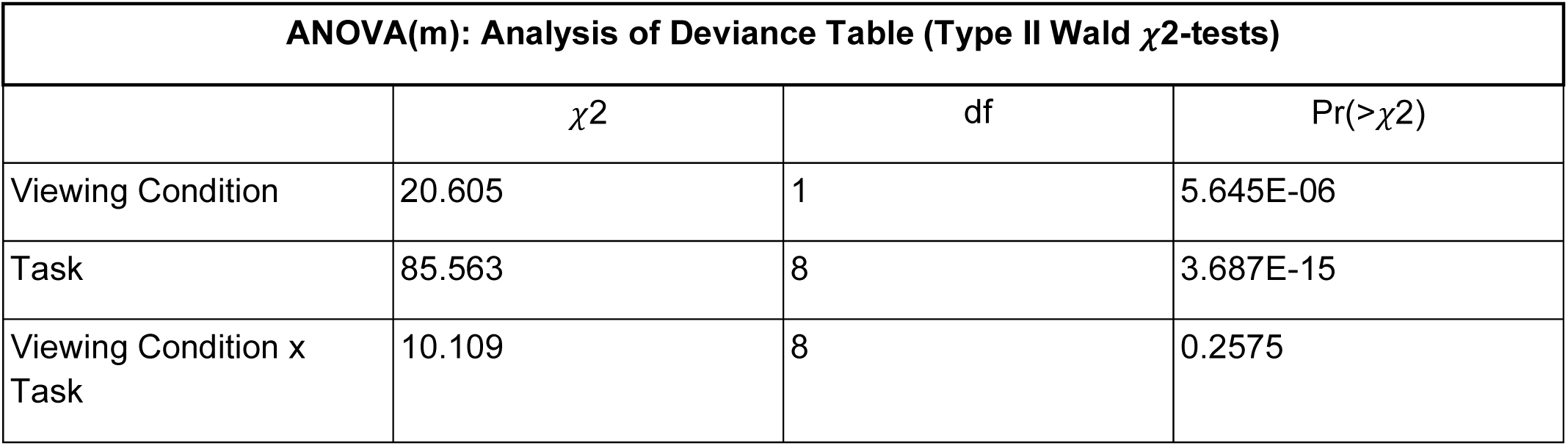
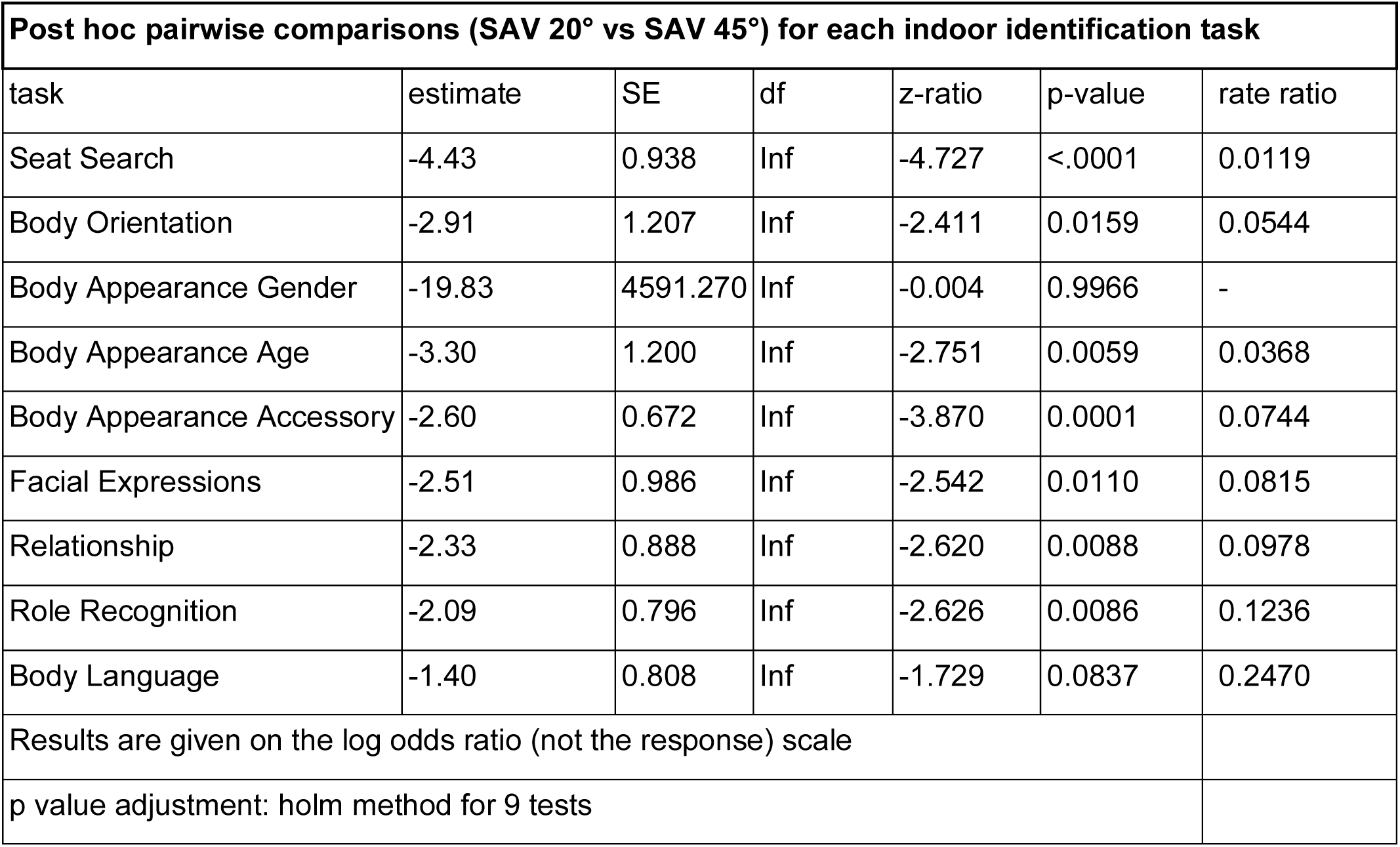

| ANOVA(m): Analysis of Deviance Table (Type II Wald $\chi^2$-tests) | | | |
| --- | --- | --- | --- |
| | $\chi^2$ | df | Pr(>$\chi^2$) |
| Viewing Condition | 20.605 | 1 | 5.645E-06 |
| Task | 85.563 | 8 | 3.687E-15 |
| Viewing Condition x Task | 10.109 | 8 | 0.2575 |

| Post hoc pairwise comparisons (SAV 20° vs SAV 45°) for each indoor identification task |  |  |  |  |  |  |
| --- | --- | --- | --- | --- | --- | --- |
| task | estimate | SE | df | z-ratio | p-value | rate ratio |
| Seat Search | -4.43 | 0.938 | Inf | -4.727 | <.0001 | 0.0119 |
| Body Orientation | -2.91 | 1.207 | Inf | -2.411 | 0.0159 | 0.0544 |
| Body Appearance Gender | -19.83 | 4591.270 | Inf | -0.004 | 0.9966 | - |
| Body Appearance Age | -3.30 | 1.200 | Inf | -2.751 | 0.0059 | 0.0368 |
| Body Appearance Accessory | -2.60 | 0.672 | Inf | -3.870 | 0.0001 | 0.0744 |
| Facial Expressions | -2.51 | 0.986 | Inf | -2.542 | 0.0110 | 0.0815 |
| Relationship | -2.33 | 0.888 | Inf | -2.620 | 0.0088 | 0.0978 |
| Role Recognition | -2.09 | 0.796 | Inf | -2.626 | 0.0086 | 0.1236 |
| Body Language | -1.40 | 0.808 | Inf | -1.729 | 0.0837 | 0.2470 |
| Results are given on the log odds ratio (not the response) scale |  |  |  |  |  |  |
| p value adjustment: holm method for 9 tests |  |  |  |  |  |  |

- Indoor identification tasks: Decision time

Best fitting model: Generalized linear mixed-effects model with a tweedie distribution. The dependent variable was the decision time, with user condition, task, and their interaction as fixed effects. Since the design involved repeated measures, participant ID was included as a random effect. A log offset for the total number of trials was applied to account for differences in trial counts across tasks.

Model assumptions:

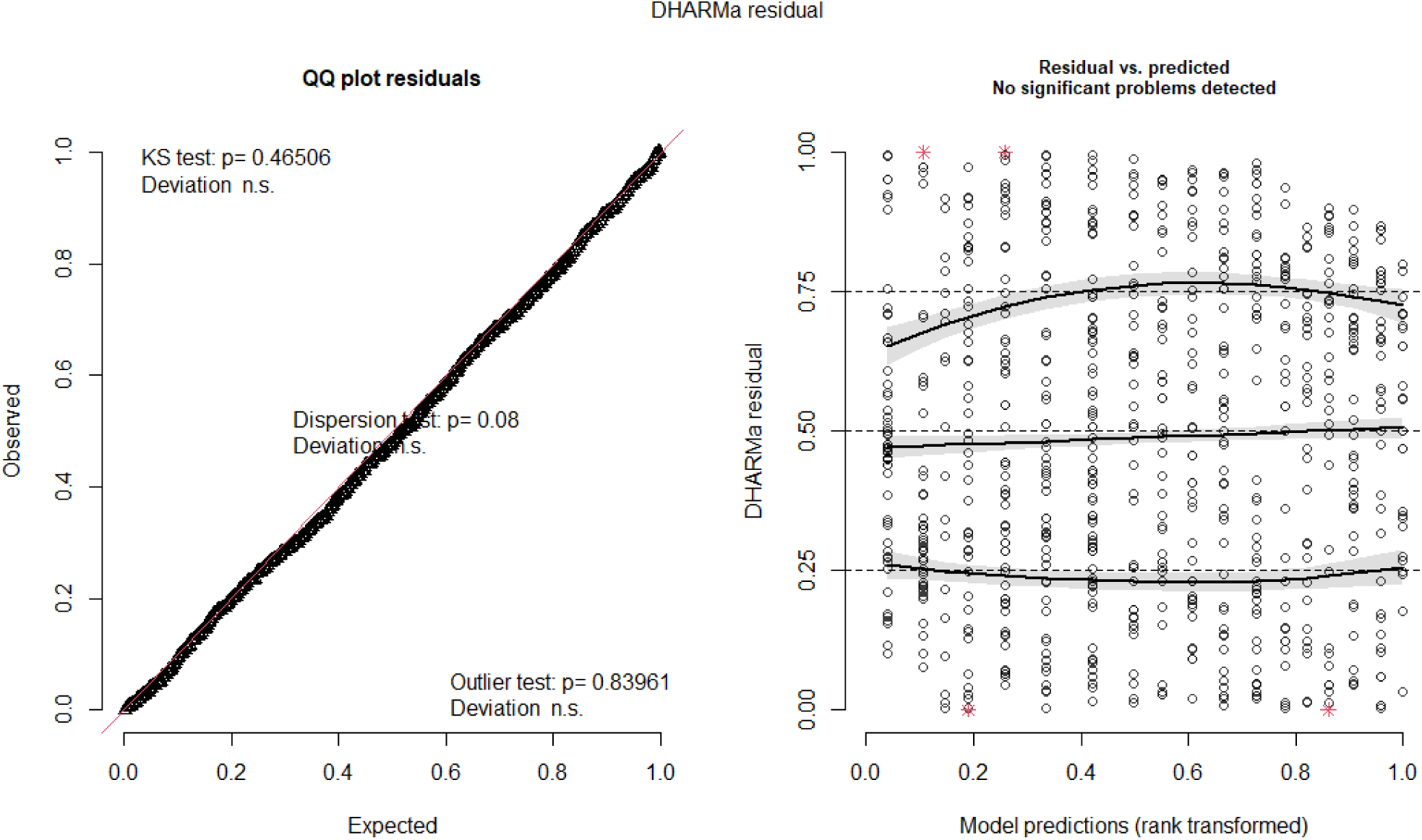

Statistical results:

| ANOVA(m): Analysis of Deviance Table (Type II Wald $\chi^2$-tests) | | | |
| --- | --- | --- | --- |
| | $\chi^2$ | df | Pr(>$\chi^2$) |
| Viewing Condition | 25.084 | 1 | 5.489E-07 |
| Task | 1485.231 | 8 | <2.2E-16 |
| Viewing Condition x Task | 37.659 | 8 | 8.703E-06 |

| Post hoc pairwise comparisons (SAV 20° vs SAV 45°) for each indoor identification task |  |  |  |  |  |  |
| --- | --- | --- | --- | --- | --- | --- |
| task | estimate | SE | df | z-ratio | p-value | rate ratio |
| Seat Search | 0.6728 | 0.113 | Inf | 5.934 | <.0001 | 1.960 |
| Body Orientation | 0.5854 | 0.113 | Inf | 5.179 | <.0001 | 1.796 |
| Body Appearance Gender | 0.4879 | 0.113 | Inf | 4.315 | <.0001 | 1.629 |
| Body Appearance Age | 0.5370 | 0.140 | Inf | 3.846 | 0.0001 | 1.711 |
| Body Appearance Accessory | -0.0372 | 0.155 | Inf | -0.240 | 0.8102 | - |
| Facial Expressions | 0.2858 | 0.125 | Inf | 2.294 | 0.0218 | 1.331 |
| Relationship | 0.2118 | 0.125 | Inf | 1.696 | 0.0899 | - |
| Role Recognition | 0.2138 | 0.124 | Inf | 1.722 | 0.0851 | - |
| Body Language | 0.2248 | 0.123 | Inf | 1.825 | 0.0680 | - |
| Results are given on the log odds ratio (not the response) scale |  |  |  |  |  |  |
| p value adjustment: holm method for 9 tests |  |  |  |  |  |  |

- Indoor identification tasks: Confidence

Best fitting model: Cumulative link mixed-effects model (CLMM) with a logit link function. Confidence ratings were the dependent variable, with user condition, task, and their interaction as fixed effects. Since the design involved repeated measures, participant ID was included as a random effect.

Statistical results:

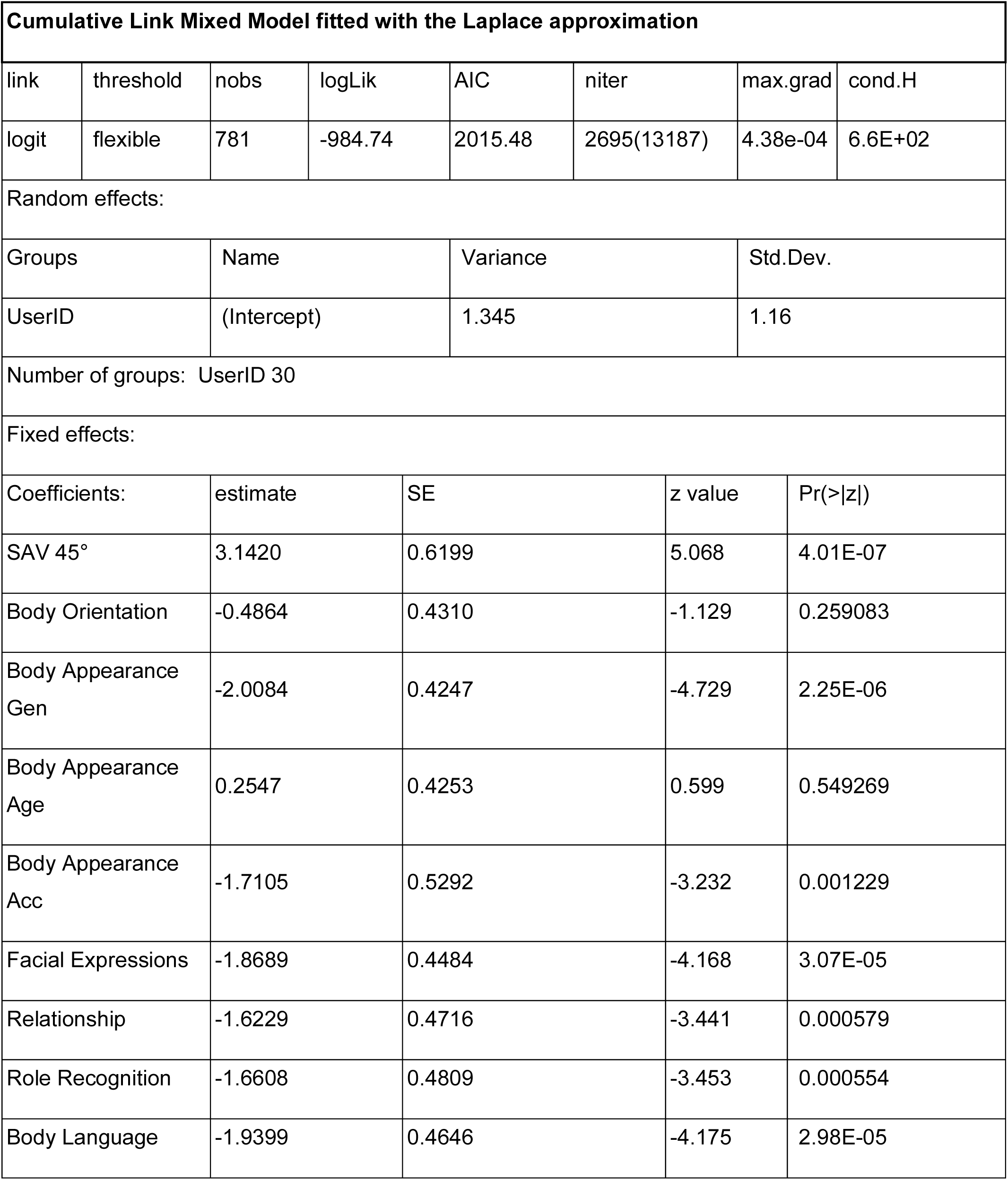
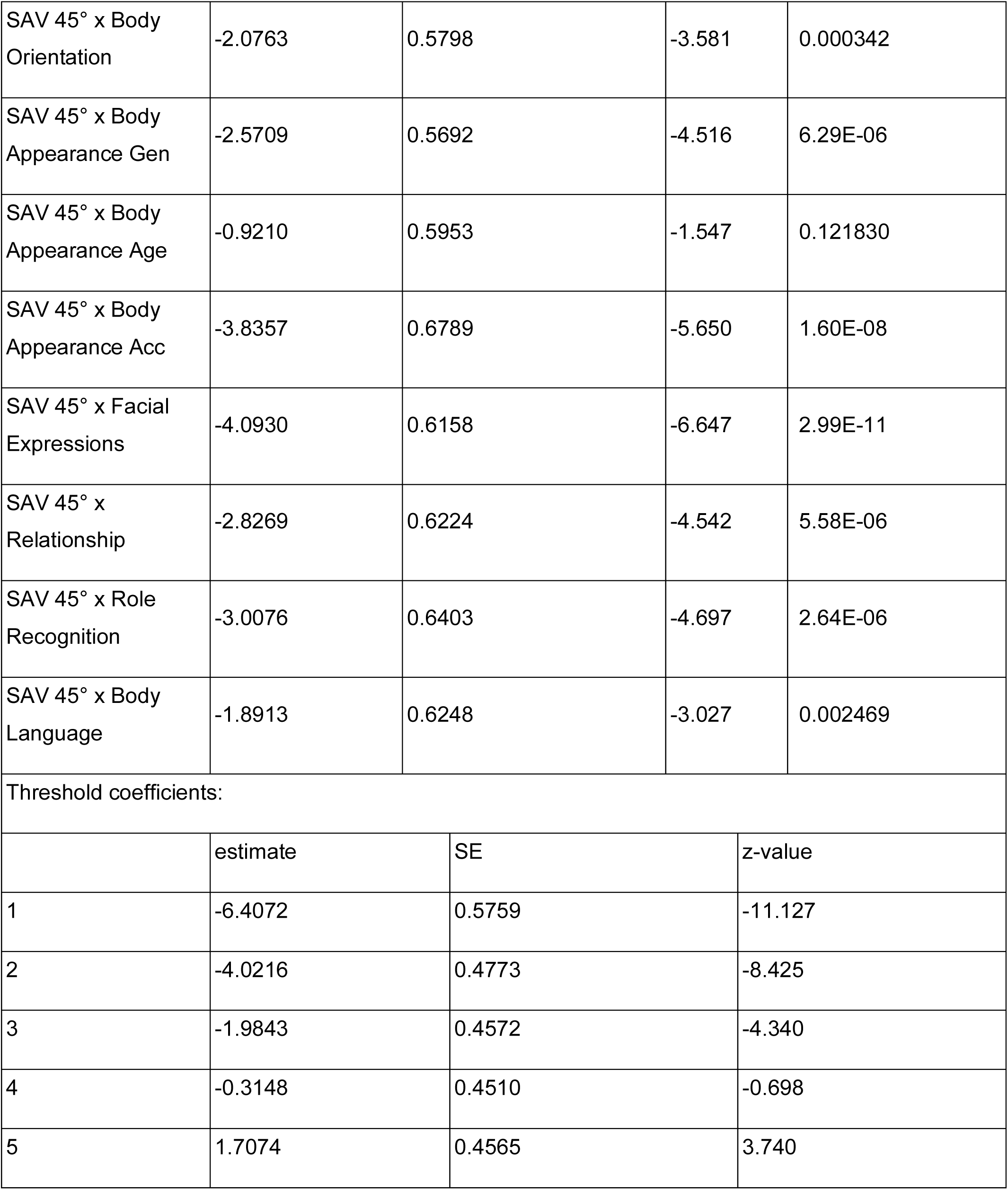
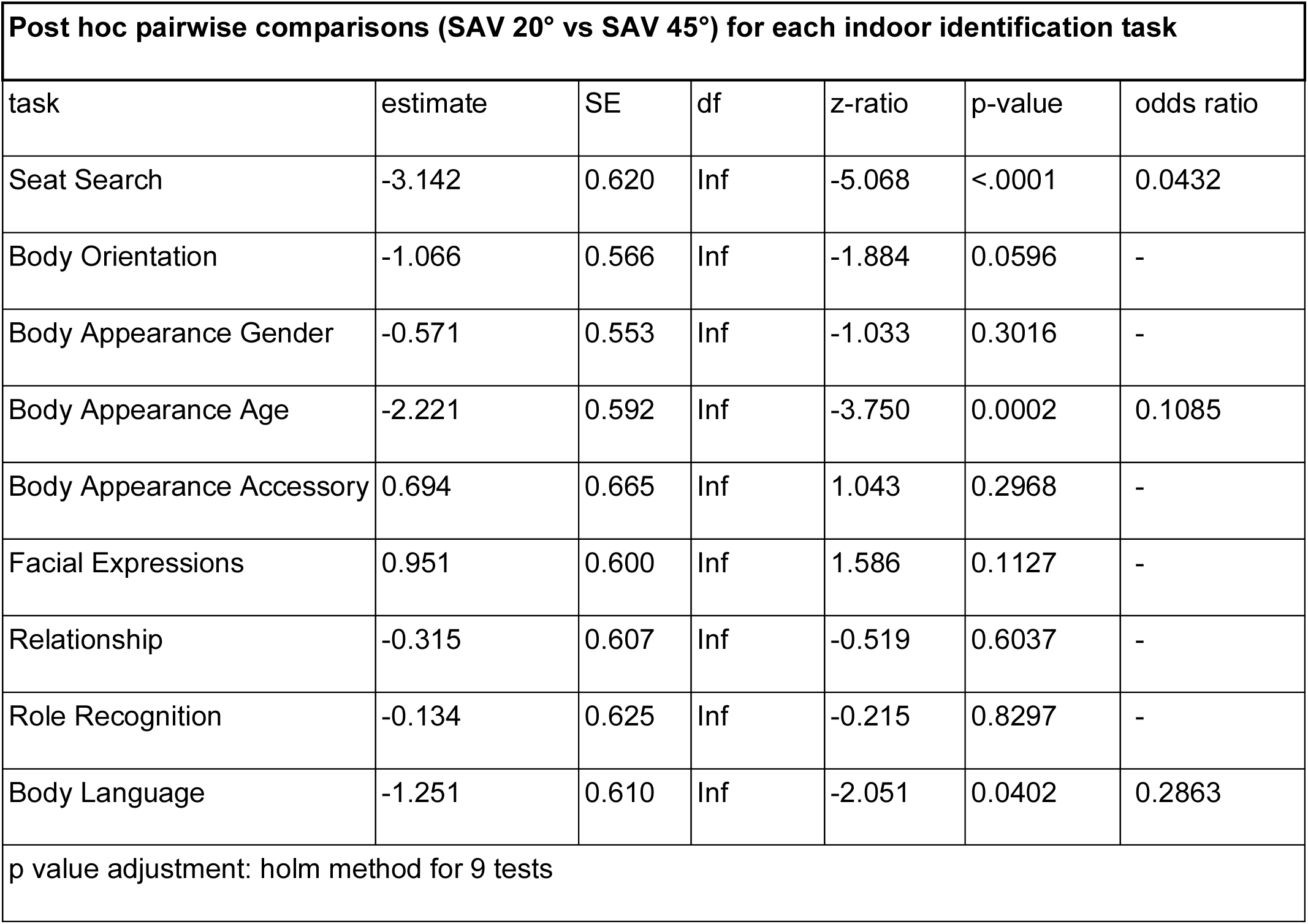

| Cumulative Link Mixed Model fitted with the Laplace approximation |  |  |  |  |  |  |  |  |
| --- | --- | --- | --- | --- | --- | --- | --- | --- |
| link | threshold | nobs | logLik | AIC | niter | max.grad | cond.H |  |
| logit | flexible | 781 | -984.74 | 2015.48 | 2695(13187) | 4.38e-04 | 6.6E+02 |  |
| Random effects: |  |  |  |  |  |  |  |  |
| Groups |  | Name |  | Variance |  | Std.Dev. |  |  |
| UserID |  | (Intercept) |  | 1.345 |  | 1.16 |  |  |
| Number of groups: UserID 30 |  |  |  |  |  |  |  |  |
| Fixed effects: |  |  |  |  |  |  |  |  |
| Coefficients: |  | estimate |  | SE |  | z value |  | Pr(>\|z\|) |
| SAV 45° |  | 3.1420 |  | 0.6199 |  | 5.068 |  | 4.01E-07 |
| Body Orientation |  | -0.4864 |  | 0.4310 |  | -1.129 |  | 0.259083 |
| Body Appearance Gen |  | -2.0084 |  | 0.4247 |  | -4.729 |  | 2.25E-06 |
| Body Appearance Age |  | 0.2547 |  | 0.4253 |  | 0.599 |  | 0.549269 |
| Body Appearance Acc |  | -1.7105 |  | 0.5292 |  | -3.232 |  | 0.001229 |
| Facial Expressions |  | -1.8689 |  | 0.4484 |  | -4.168 |  | 3.07E-05 |
| Relationship |  | -1.6229 |  | 0.4716 |  | -3.441 |  | 0.000579 |
| Role Recognition |  | -1.6608 |  | 0.4809 |  | -3.453 |  | 0.000554 |
| Body Language |  | -1.9399 |  | 0.4646 |  | -4.175 |  | 2.98E-05 |

|  |  |  |  |  |
| --- | --- | --- | --- | --- |
| SAV 45° x Body Orientation | -2.0763 | 0.5798 | -3.581 | 0.000342 |
| SAV 45° x Body Appearance Gen | -2.5709 | 0.5692 | -4.516 | 6.29E-06 |
| SAV 45° x Body Appearance Age | -0.9210 | 0.5953 | -1.547 | 0.121830 |
| SAV 45° x Body Appearance Acc | -3.8357 | 0.6789 | -5.650 | 1.60E-08 |
| SAV 45° x Facial Expressions | -4.0930 | 0.6158 | -6.647 | 2.99E-11 |
| SAV 45° x Relationship | -2.8269 | 0.6224 | -4.542 | 5.58E-06 |
| SAV 45° x Role Recognition | -3.0076 | 0.6403 | -4.697 | 2.64E-06 |
| SAV 45° x Body Language | -1.8913 | 0.6248 | -3.027 | 0.002469 |
| Threshold coefficients: |  |  |  |  |
|  | estimate | SE | z-value |  |
| 1 | -6.4072 | 0.5759 | -11.127 |  |
| 2 | -4.0216 | 0.4773 | -8.425 |  |
| 3 | -1.9843 | 0.4572 | -4.340 |  |
| 4 | -0.3148 | 0.4510 | -0.698 |  |
| 5 | 1.7074 | 0.4565 | 3.740 |  |

| Post hoc pairwise comparisons (SAV 20° vs SAV 45°) for each indoor identification task |  |  |  |  |  |  |
| --- | --- | --- | --- | --- | --- | --- |
| task | estimate | SE | df | z-ratio | p-value | odds ratio |
| Seat Search | -3.142 | 0.620 | Inf | -5.068 | <.0001 | 0.0432 |
| Body Orientation | -1.066 | 0.566 | Inf | -1.884 | 0.0596 | - |
| Body Appearance Gender | -0.571 | 0.553 | Inf | -1.033 | 0.3016 | - |
| Body Appearance Age | -2.221 | 0.592 | Inf | -3.750 | 0.0002 | 0.1085 |
| Body Appearance Accessory | 0.694 | 0.665 | Inf | 1.043 | 0.2968 | - |
| Facial Expressions | 0.951 | 0.600 | Inf | 1.586 | 0.1127 | - |
| Relationship | -0.315 | 0.607 | Inf | -0.519 | 0.6037 | - |
| Role Recognition | -0.134 | 0.625 | Inf | -0.215 | 0.8297 | - |
| Body Language | -1.251 | 0.610 | Inf | -2.051 | 0.0402 | 0.2863 |
| p value adjustment: holm method for 9 tests |  |  |  |  |  |  |

- Indoor identification tasks: Success rate

Best fitting model: Generalized linear mixed-effects model with a binomial distribution. Success rate was the dependent variable, with viewing condition, task, and their interaction as fixed effects. Since the design follows a repeated measures structure, participant was included as a random effect. Additionally, a log-transformed total trial count was used as an offset to account for differences in trial numbers across conditions.

Model assumptions:

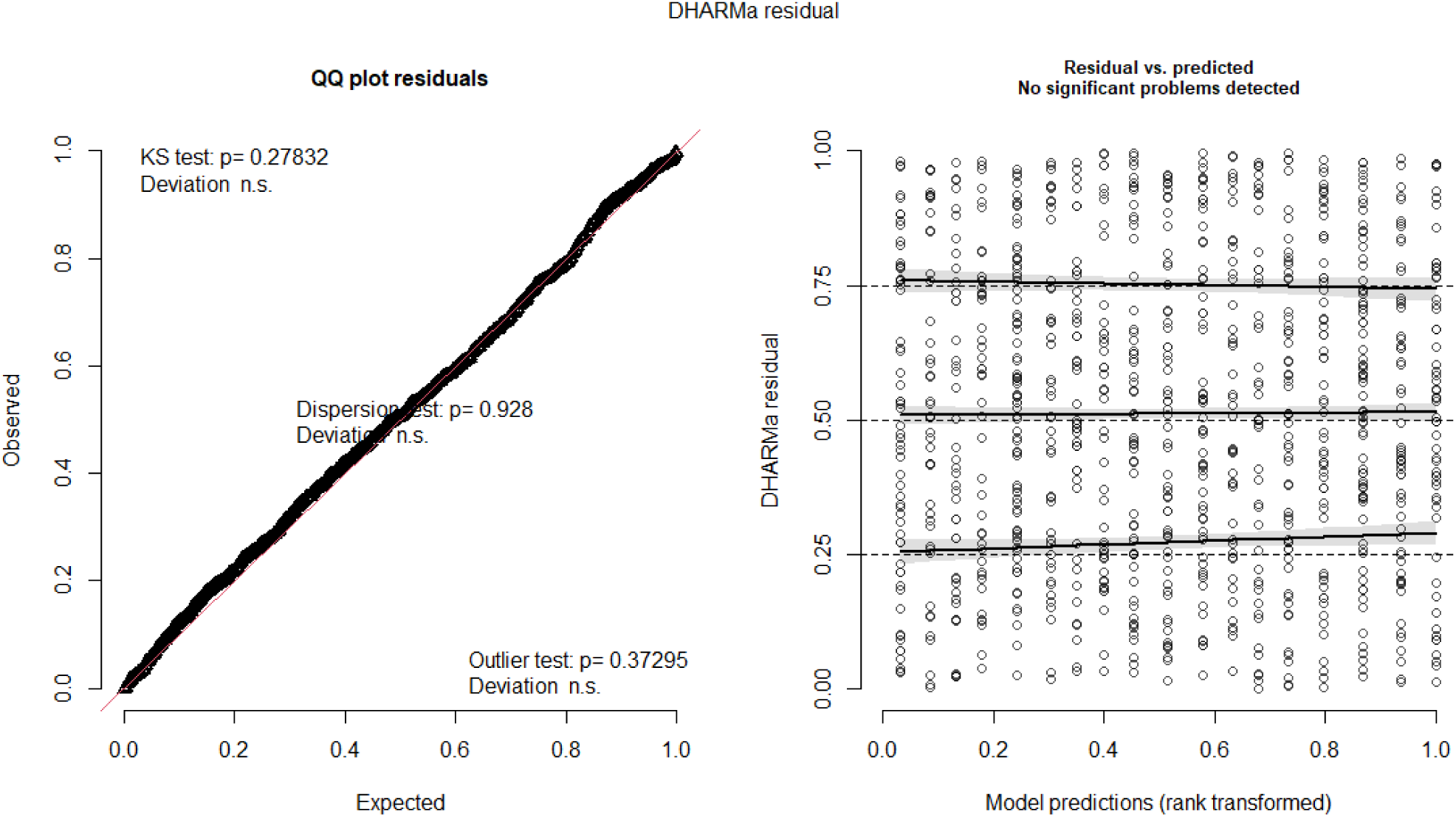

Statistical results:

| ANOVA(m): Analysis of Deviance Table (Type II Wald $\chi^2$-tests) | | | |
| --- | --- | --- | --- |
| | $\chi^2$ | df | Pr(>$\chi^2$) |
| Viewing Condition | 18.693 | 1 | 1.535E-05 |
| Task | 79.596 | 8 | 5.895E-14 |
| Viewing Condition x Task | 44.063 | 8 | 5.535E-07 |

| Post hoc pairwise comparisons (SAV 20° vs SAV 45°) for each indoor identification task |  |  |  |  |  |  |
| --- | --- | --- | --- | --- | --- | --- |
| task | estimate | SE | df | z-ratio | p-value | rate ratio |
| Seat Search | -3.1737 | 0.562 | Inf | -5.647 | <.0001 | 0.0418 |
| Body Orientation | -2.5531 | 0.645 | Inf | -3.961 | 0.0001 | 0.0778 |
| Body Appearance Gender | -1.7231 | 0.516 | Inf | -3.342 | 0.0008 | 0.1785 |
| Body Appearance Age | -2.2098 | 0.714 | Inf | -3.097 | 0.0020 | 0.1097 |
| Body Appearance Accessory | -1.5368 | 0.506 | Inf | -3.034 | 0.0024 | 0.2151 |
| Facial Expressions | 0.2710 | 0.536 | Inf | 0.506 | 0.6131 | - |
| Relationship | 0.0723 | 0.511 | Inf | 0.141 | 0.8876 | - |
| Role Recognition | -1.5678 | 0.534 | Inf | -2.934 | 0.0033 | 0.2085 |
| Body Language | -0.6122 | 0.513 | Inf | -1.193 | 0.2329 | - |
| Results are given on the log odds ratio (not the response) scale |  |  |  |  |  |  |
| p value adjustment: holm method for 9 tests |  |  |  |  |  |  |

- Indoor identification tasks: Decision time when successful

Best fitting model: Generalized linear mixed-effects model with a Tweedie distribution. Decision duration was the dependent variable, modeled only for successful trials. Viewing condition, task, and their interaction were included as fixed effects, while participant was included as a random effect to account for the repeated measures design. A log-transformed total trial count was used as an offset to control for differences in trial numbers across conditions. Additionally, dispersion was modeled as a function of the viewing condition to account for potential heterogeneity in variance.

Model assumptions:

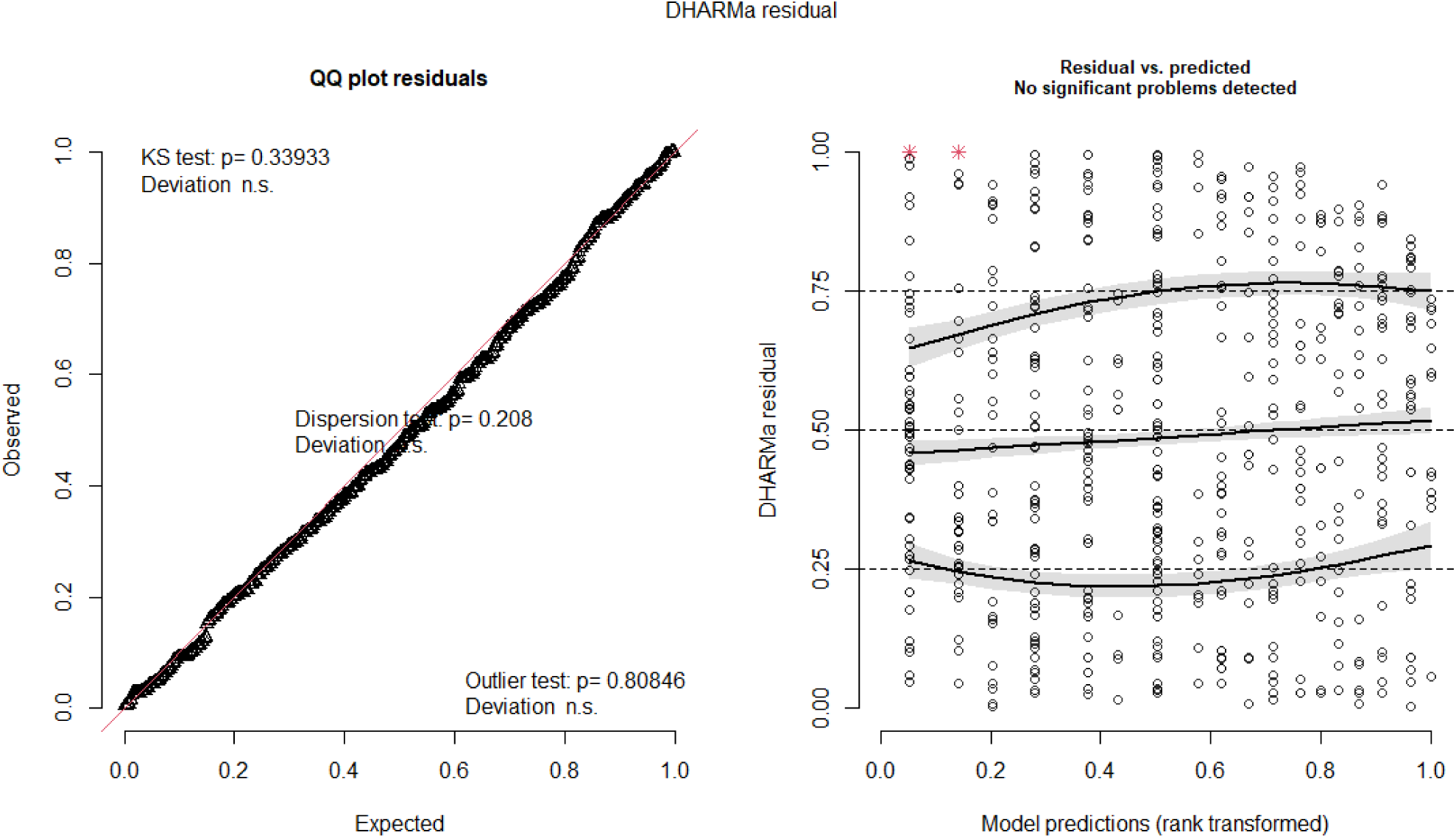

Statistical results:

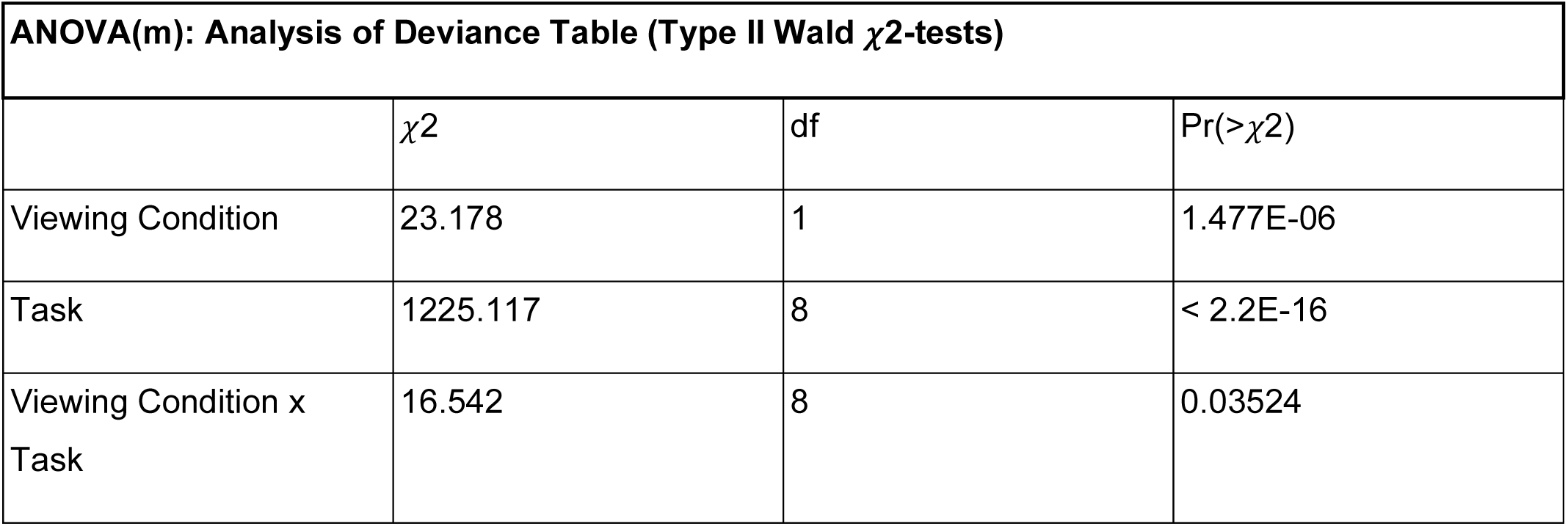
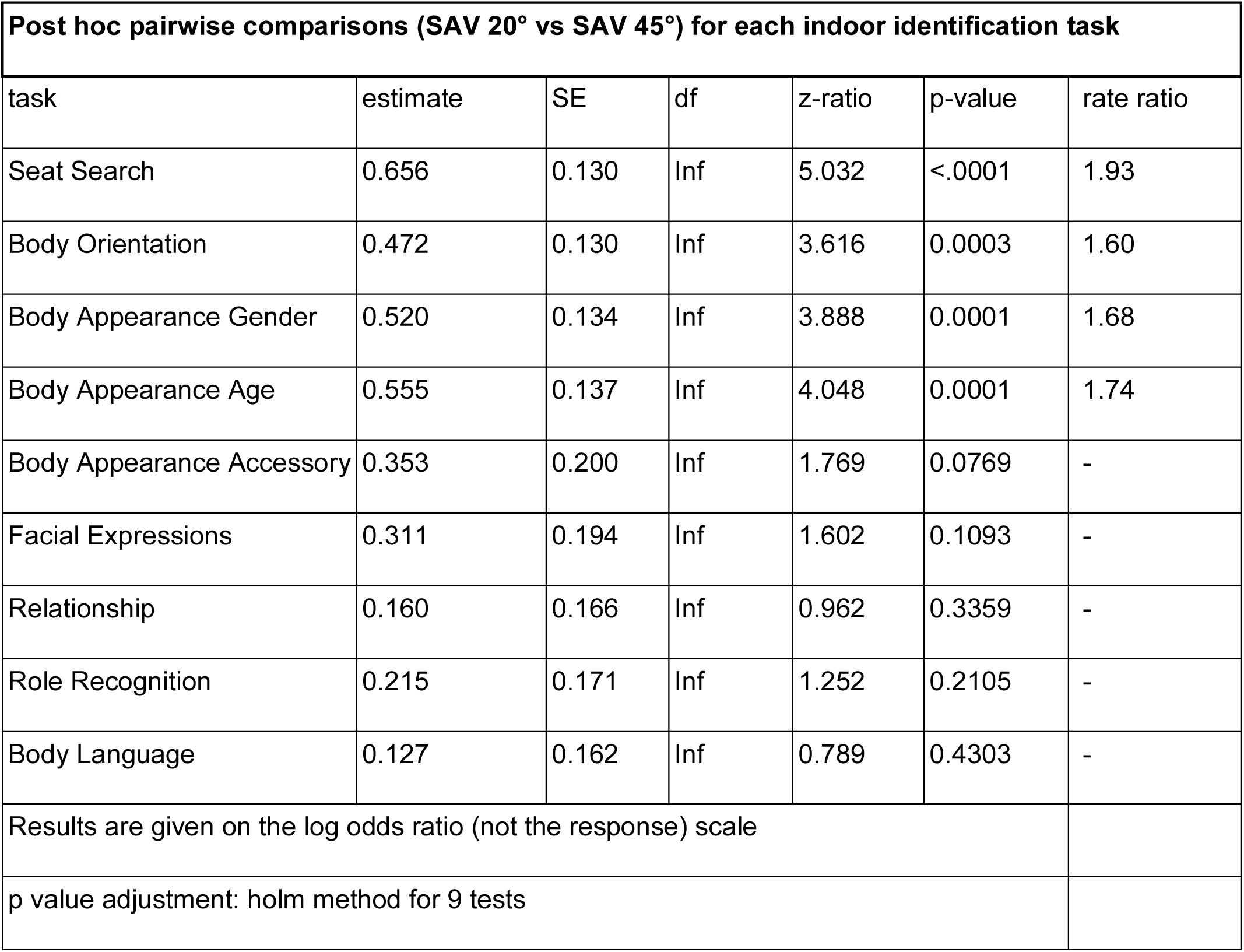

| ANOVA(m): Analysis of Deviance Table (Type II Wald $\chi^2$-tests) | | | |
| --- | --- | --- | --- |
| | $\chi^2$ | df | Pr(>$\chi^2$) |
| Viewing Condition | 23.178 | 1 | 1.477E-06 |
| Task | 1225.117 | 8 | < 2.2E-16 |
| Viewing Condition x Task | 16.542 | 8 | 0.03524 |

| Post hoc pairwise comparisons (SAV 20° vs SAV 45°) for each indoor identification task |  |  |  |  |  |  |
| --- | --- | --- | --- | --- | --- | --- |
| task | estimate | SE | df | z-ratio | p-value | rate ratio |
| Seat Search | 0.656 | 0.130 | Inf | 5.032 | <.0001 | 1.93 |
| Body Orientation | 0.472 | 0.130 | Inf | 3.616 | 0.0003 | 1.60 |
| Body Appearance Gender | 0.520 | 0.134 | Inf | 3.888 | 0.0001 | 1.68 |
| Body Appearance Age | 0.555 | 0.137 | Inf | 4.048 | 0.0001 | 1.74 |
| Body Appearance Accessory | 0.353 | 0.200 | Inf | 1.769 | 0.0769 | - |
| Facial Expressions | 0.311 | 0.194 | Inf | 1.602 | 0.1093 | - |
| Relationship | 0.160 | 0.166 | Inf | 0.962 | 0.3359 | - |
| Role Recognition | 0.215 | 0.171 | Inf | 1.252 | 0.2105 | - |
| Body Language | 0.127 | 0.162 | Inf | 0.789 | 0.4303 | - |
| Results are given on the log odds ratio (not the response) scale |  |  |  |  |  |  |
| p value adjustment: holm method for 9 tests |  |  |  |  |  |  |

- Indoor identification tasks: Confidence when successful

Best fitting model: Cumulative link mixed-effects model (CLMM) with a logit link function. Confidence ratings for successful trials were the dependent variable, with user condition, task, and their interaction as fixed effects. Since the design involved repeated measures, participant ID was included as a random effect.

Statistical results:

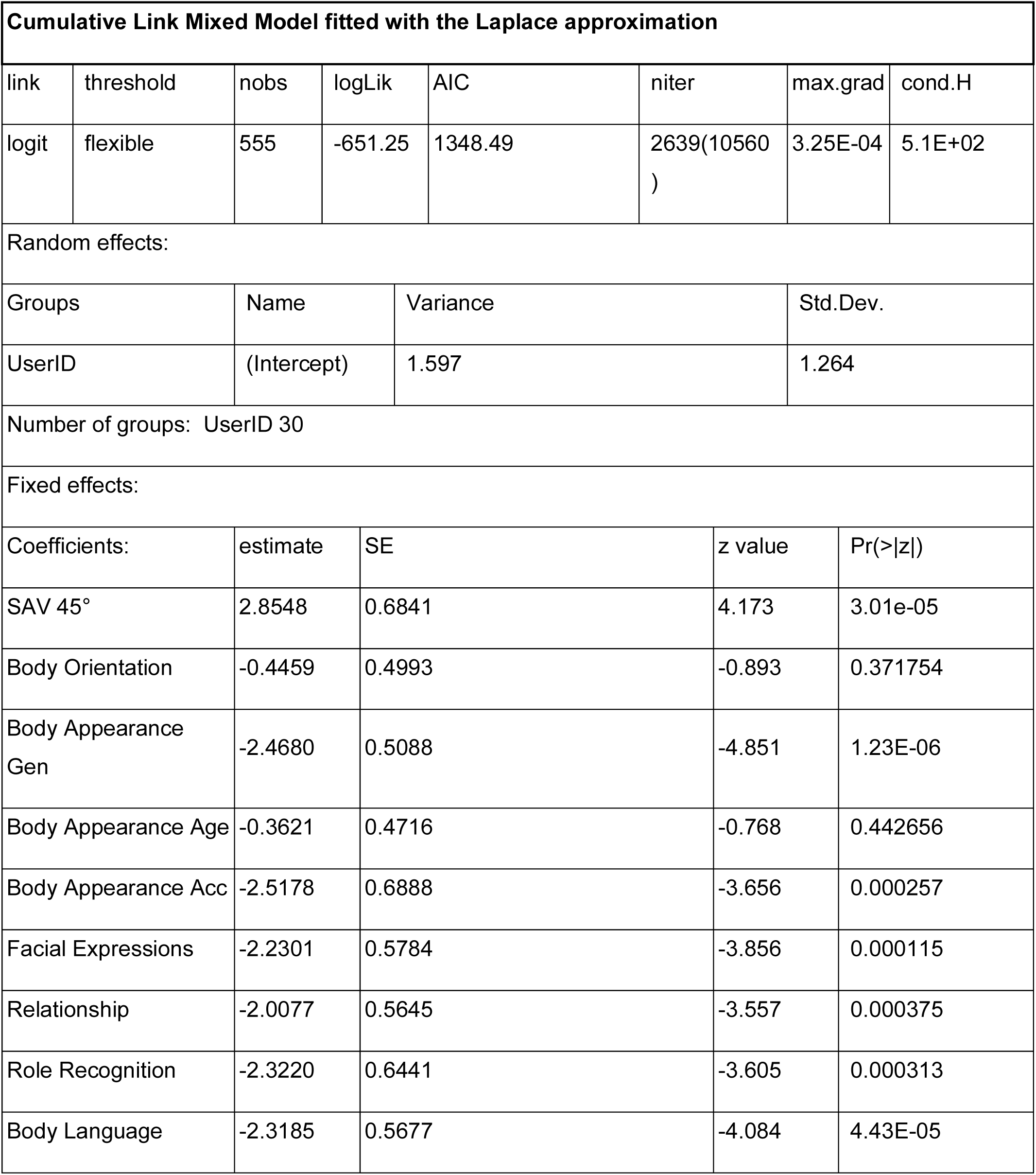
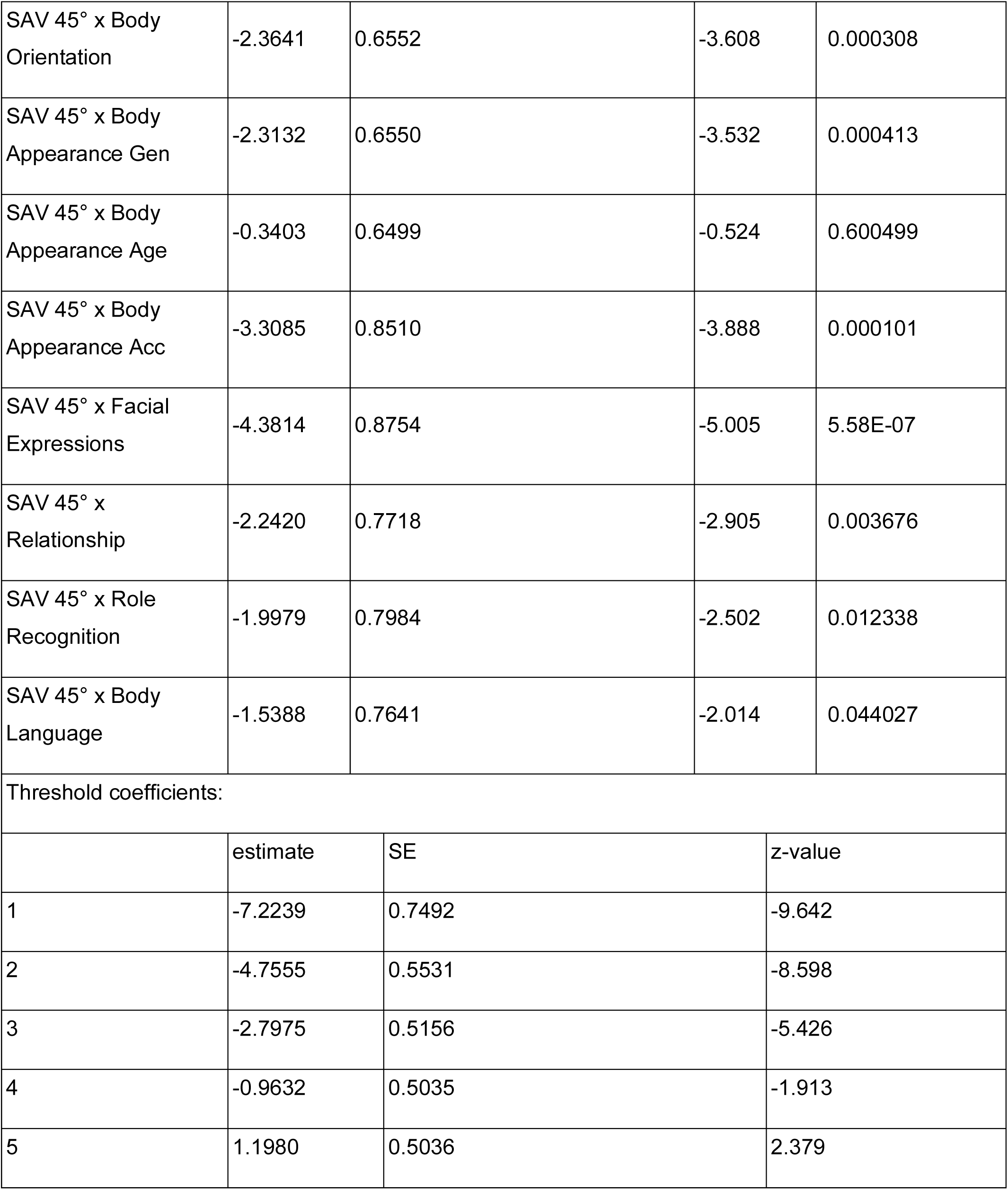
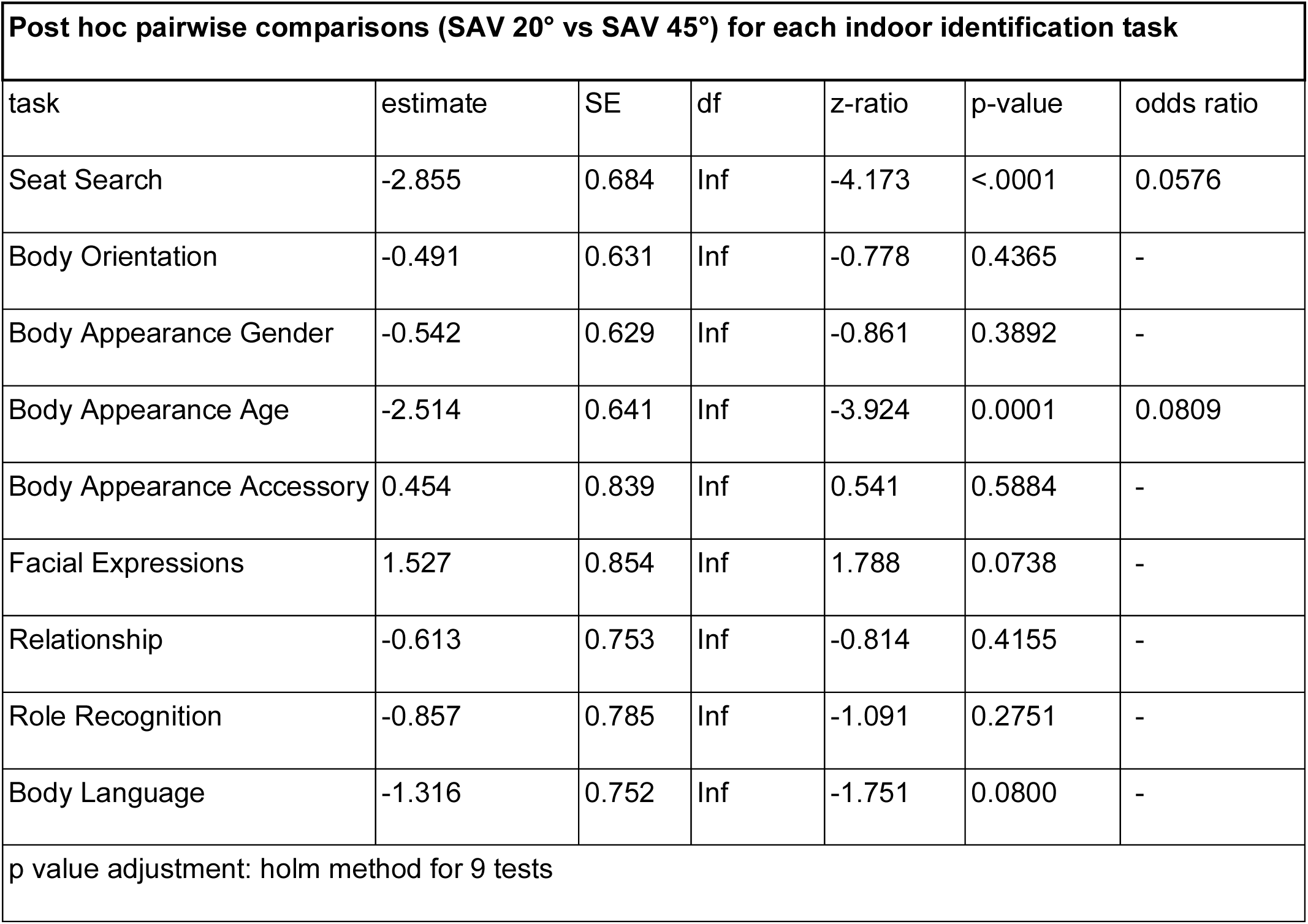

| Cumulative Link Mixed Model fitted with the Laplace approximation |  |  |  |  |  |  |  |
| --- | --- | --- | --- | --- | --- | --- | --- |
| link | threshold | nobs | logLik | AIC | niter | max.grad | cond.H |
| logit | flexible | 555 | -651.25 | 1348.49 | 2639(10560<br>) | 3.25E-04 | 5.1E+02 |
| Random effects: |  |  |  |  |  |  |  |
| Groups |  | Name | Variance |  |  | Std.Dev. |  |
| UserID |  | (Intercept) | 1.597 |  |  | 1.264 |  |
| Number of groups: UserID 30 |  |  |  |  |  |  |  |
| Fixed effects: |  |  |  |  |  |  |  |
| Coefficients: |  | estimate | SE |  | z value | Pr(>\|z\|) |  |
| SAV 45° |  | 2.8548 | 0.6841 |  | 4.173 | 3.01e-05 |  |
| Body Orientation |  | -0.4459 | 0.4993 |  | -0.893 | 0.371754 |  |
| Body Appearance Gen |  | -2.4680 | 0.5088 |  | -4.851 | 1.23E-06 |  |
| Body Appearance Age |  | -0.3621 | 0.4716 |  | -0.768 | 0.442656 |  |
| Body Appearance Acc |  | -2.5178 | 0.6888 |  | -3.656 | 0.000257 |  |
| Facial Expressions |  | -2.2301 | 0.5784 |  | -3.856 | 0.000115 |  |
| Relationship |  | -2.0077 | 0.5645 |  | -3.557 | 0.000375 |  |
| Role Recognition |  | -2.3220 | 0.6441 |  | -3.605 | 0.000313 |  |
| Body Language |  | -2.3185 | 0.5677 |  | -4.084 | 4.43E-05 |  |

|  |  |  |  |  |
| --- | --- | --- | --- | --- |
| SAV 45° x Body Orientation | -2.3641 | 0.6552 | -3.608 | 0.000308 |
| SAV 45° x Body Appearance Gen | -2.3132 | 0.6550 | -3.532 | 0.000413 |
| SAV 45° x Body Appearance Age | -0.3403 | 0.6499 | -0.524 | 0.600499 |
| SAV 45° x Body Appearance Acc | -3.3085 | 0.8510 | -3.888 | 0.000101 |
| SAV 45° x Facial Expressions | -4.3814 | 0.8754 | -5.005 | 5.58E-07 |
| SAV 45° x Relationship | -2.2420 | 0.7718 | -2.905 | 0.003676 |
| SAV 45° x Role Recognition | -1.9979 | 0.7984 | -2.502 | 0.012338 |
| SAV 45° x Body Language | -1.5388 | 0.7641 | -2.014 | 0.044027 |
| Threshold coefficients: |  |  |  |  |
|  | estimate | SE | z-value |  |
| 1 | -7.2239 | 0.7492 | -9.642 |  |
| 2 | -4.7555 | 0.5531 | -8.598 |  |
| 3 | -2.7975 | 0.5156 | -5.426 |  |
| 4 | -0.9632 | 0.5035 | -1.913 |  |
| 5 | 1.1980 | 0.5036 | 2.379 |  |

| Post hoc pairwise comparisons (SAV 20° vs SAV 45°) for each indoor identification task |  |  |  |  |  |  |
| --- | --- | --- | --- | --- | --- | --- |
| task | estimate | SE | df | z-ratio | p-value | odds ratio |
| Seat Search | -2.855 | 0.684 | Inf | -4.173 | <.0001 | 0.0576 |
| Body Orientation | -0.491 | 0.631 | Inf | -0.778 | 0.4365 | - |
| Body Appearance Gender | -0.542 | 0.629 | Inf | -0.861 | 0.3892 | - |
| Body Appearance Age | -2.514 | 0.641 | Inf | -3.924 | 0.0001 | 0.0809 |
| Body Appearance Accessory | 0.454 | 0.839 | Inf | 0.541 | 0.5884 | - |
| Facial Expressions | 1.527 | 0.854 | Inf | 1.788 | 0.0738 | - |
| Relationship | -0.613 | 0.753 | Inf | -0.814 | 0.4155 | - |
| Role Recognition | -0.857 | 0.785 | Inf | -1.091 | 0.2751 | - |
| Body Language | -1.316 | 0.752 | Inf | -1.751 | 0.0800 | - |
| p value adjustment: holm method for 9 tests |  |  |  |  |  |  |

- Locomotion task: Success rate

Best fitting model: Generalized linear mixed-effects model with a binomial distribution. Locomotion success was modeled as the dependent variable, with viewing condition as a fixed effect. Since the design involved repeated measures, participant was included as a random effect to account for inter-individual variability. A dispersion parameter was also estimated to adjust for potential overdispersion in the data.

Model assumptions:

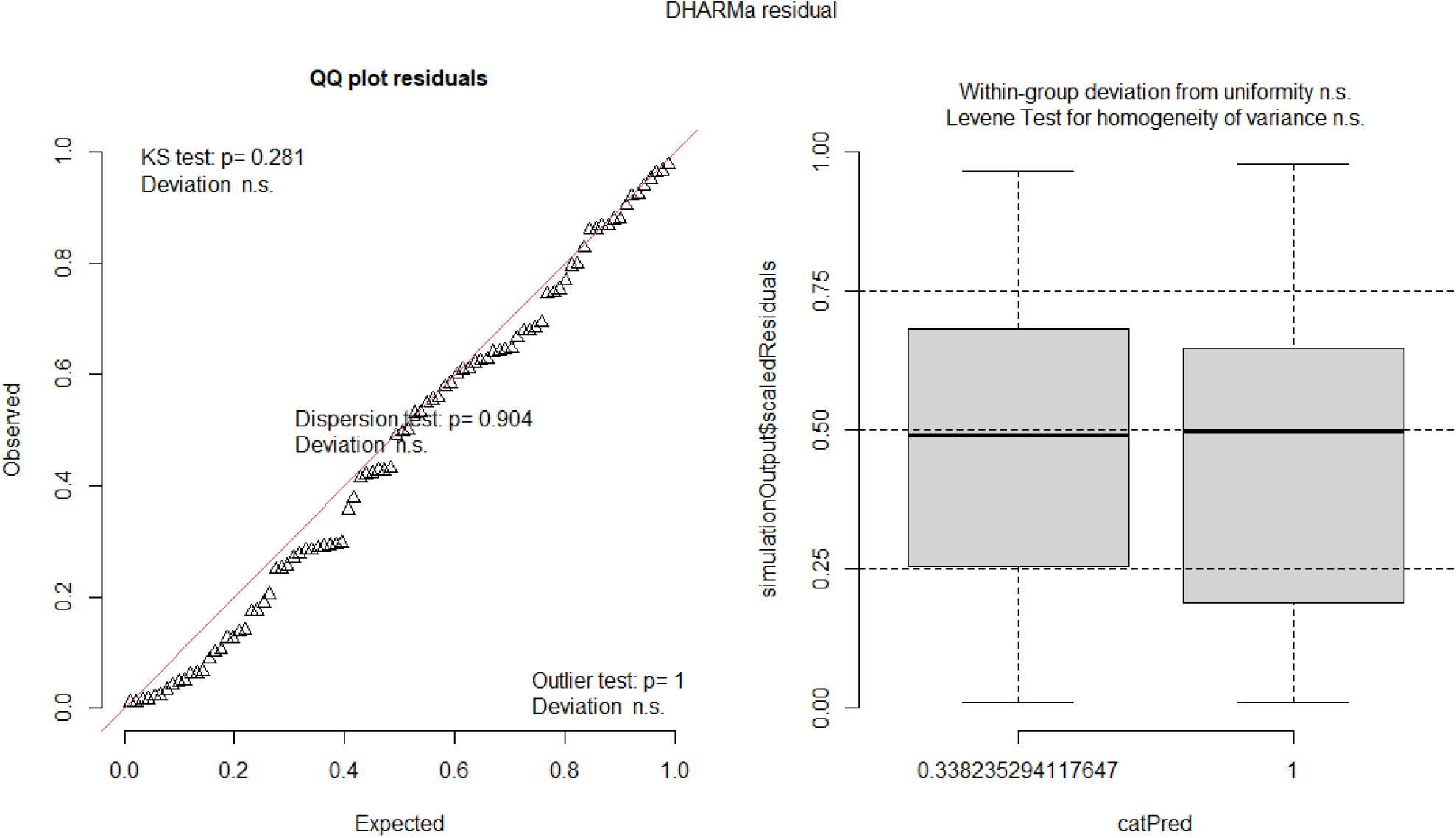

Statistical results:

| Summary of general linear mixed-effects model with binomial distribution |  |  |  |  |
| --- | --- | --- | --- | --- |
| AIC | BIC | logLik | deviance | df.resid |
| 76.0 | 83.5 | -35.0 | 70.0 | 87 |
| Random effects: |  |  |  |  |
| Groups | Name | Variance | Std.Dev. |  |
| UserID | (Intercept) | 0.2584 | 0.5083 |  |
| Number of obs: 90, groups: UserID, 30 |  |  |  |  |
| Fixed effects: |  |  |  |  |

| Coefficients: | estimate | SE | z value | Pr(>\|z\|) | Rate ratio |
| --- | --- | --- | --- | --- | --- |
| (Intercept) | -0.4318 | 0.3482 | -1.240 | 0.214920 | - |
| SAV 45° | 4.3338 | 1.1491 | 3.771 | 0.000162 | 0.0131 |
| Results are given on the log odds ratio (not the response) scale. |  |  |  |  |  |

- Locomotion task: Time in wall

Best fitting model: Generalized linear mixed-effects model with zero-inflation and dispersion correction. Time spent in the wall in the locomotion task was modeled as the dependent variable, with viewing condition as a fixed effect. Since the design involved repeated measures, participant was included as a random effect. A zero-inflation component was specified to account for excess zero values, and dispersion was modeled as a function of the viewing condition to account for variability in response distribution.

Model assumptions:

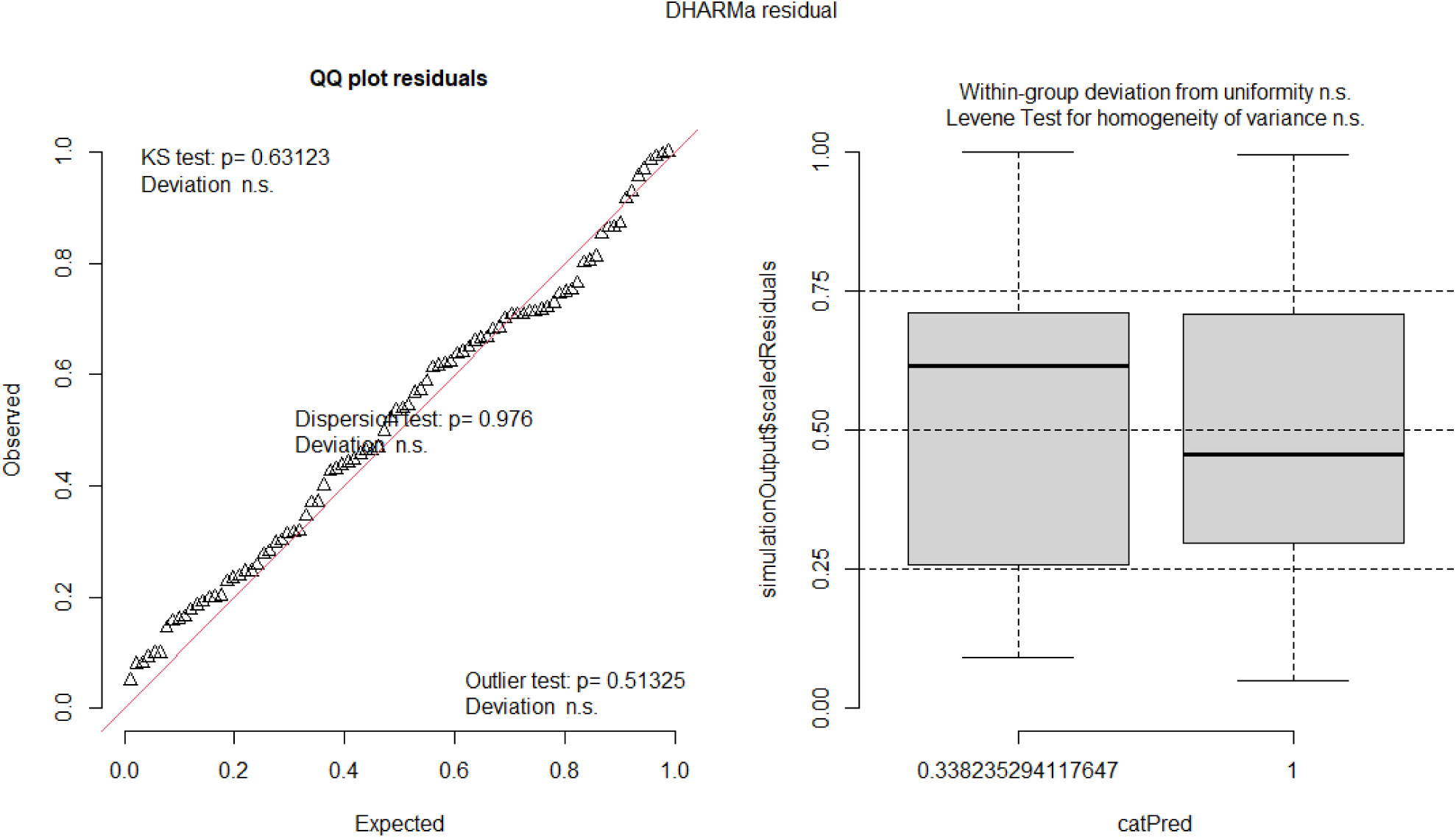

Statistical results:

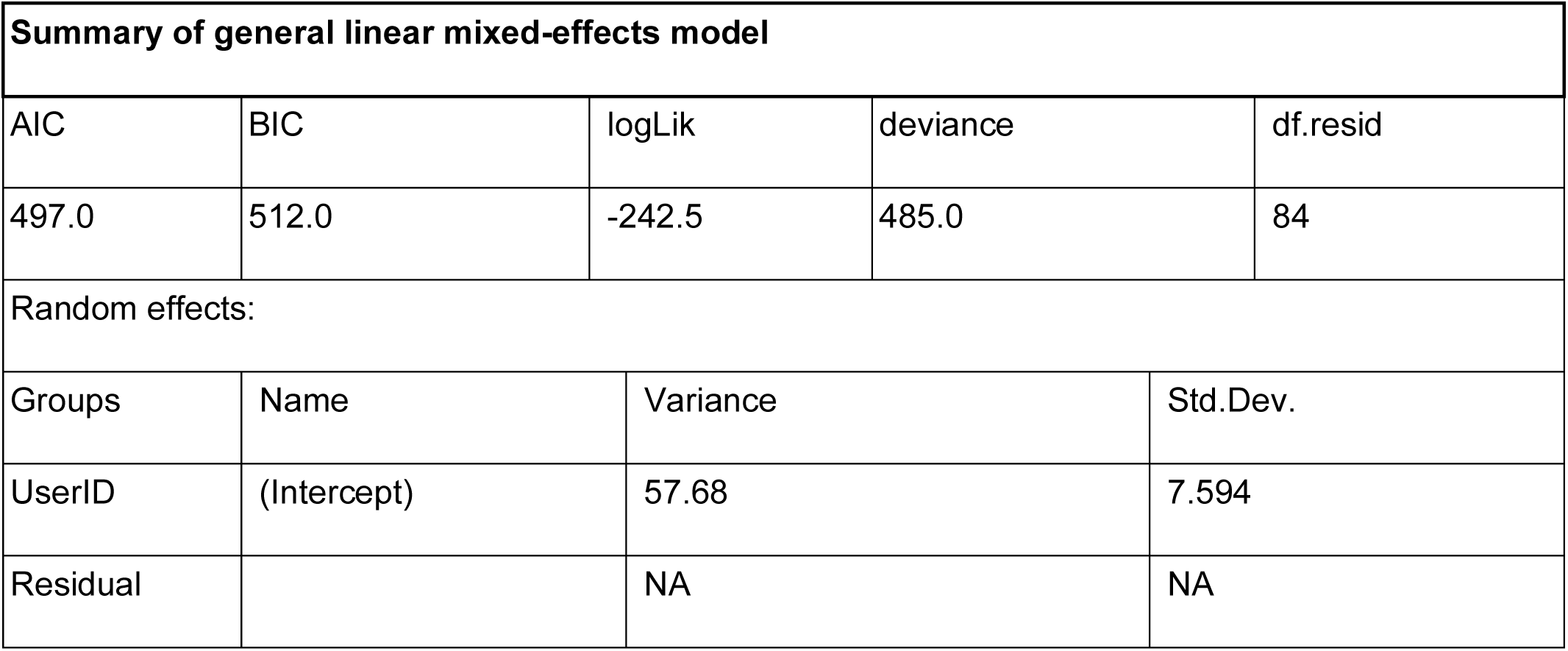
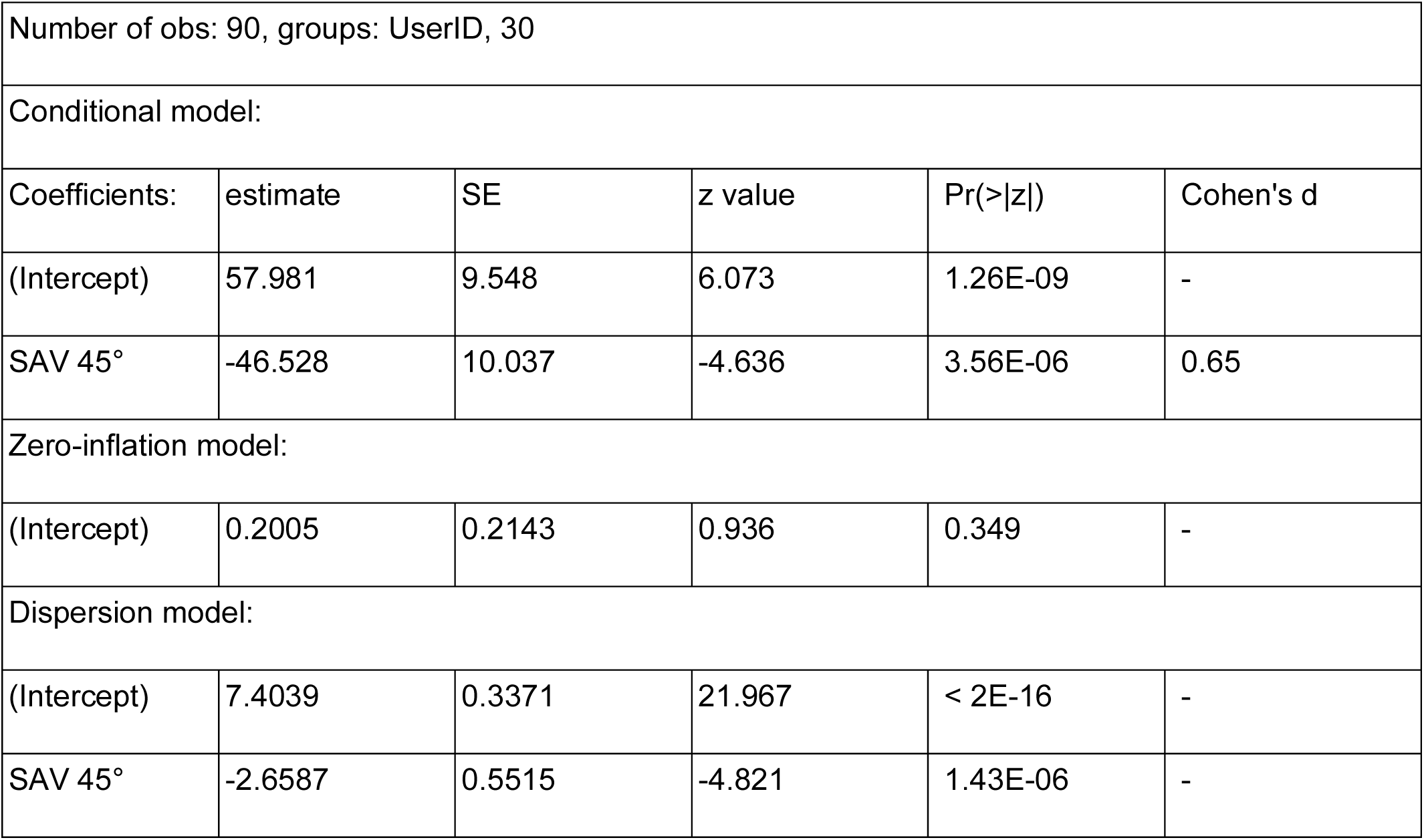

| Summary of general linear mixed-effects model |  |  |  |  |
| --- | --- | --- | --- | --- |
| AIC | BIC | logLik | deviance | df.resid |
| 497.0 | 512.0 | -242.5 | 485.0 | 84 |
| Random effects: |  |  |  |  |
| Groups | Name | Variance | Std.Dev. |  |
| UserID | (Intercept) | 57.68 | 7.594 |  |
| Residual |  | NA | NA |  |

|  |  |  |  |  |  |
| --- | --- | --- | --- | --- | --- |
| Number of obs: 90, groups: UserID, 30 |  |  |  |  |  |
| Conditional model: |  |  |  |  |  |
| Coefficients: | estimate | SE | z value | Pr(>\|z\|) | Cohen's d |
| (Intercept) | 57.981 | 9.548 | 6.073 | 1.26E-09 | - |
| SAV 45° | -46.528 | 10.037 | -4.636 | 3.56E-06 | 0.65 |
| Zero-inflation model: |  |  |  |  |  |
| (Intercept) | 0.2005 | 0.2143 | 0.936 | 0.349 | - |
| Dispersion model: |  |  |  |  |  |
| (Intercept) | 7.4039 | 0.3371 | 21.967 | < 2E-16 | - |
| SAV 45° | -2.6587 | 0.5515 | -4.821 | 1.43E-06 | - |

- Locomotion task: Mean distance

Best fitting model: Generalized linear mixed-effects model with zero-inflation and dispersion correction. Mean distance in the locomotion task was modeled as the dependent variable, with viewing condition as a fixed effect. Since the design involved repeated measures, participant was included as a random effect. A zero-inflation component was specified to account for participant-specific excess zero values, and dispersion was modeled as a function of the viewing condition to account for variability in response distribution.

Model assumptions:

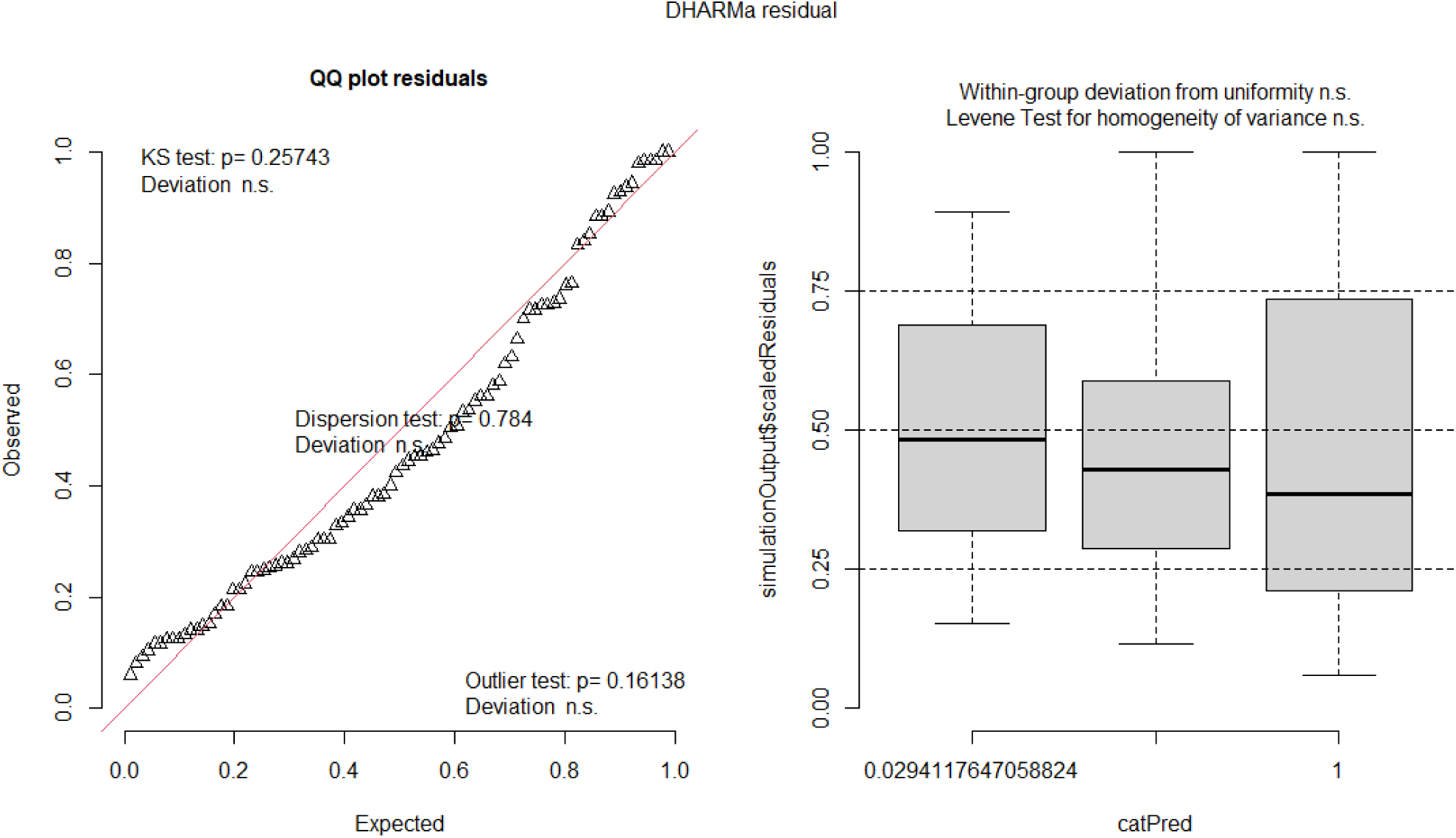

Statistical results:

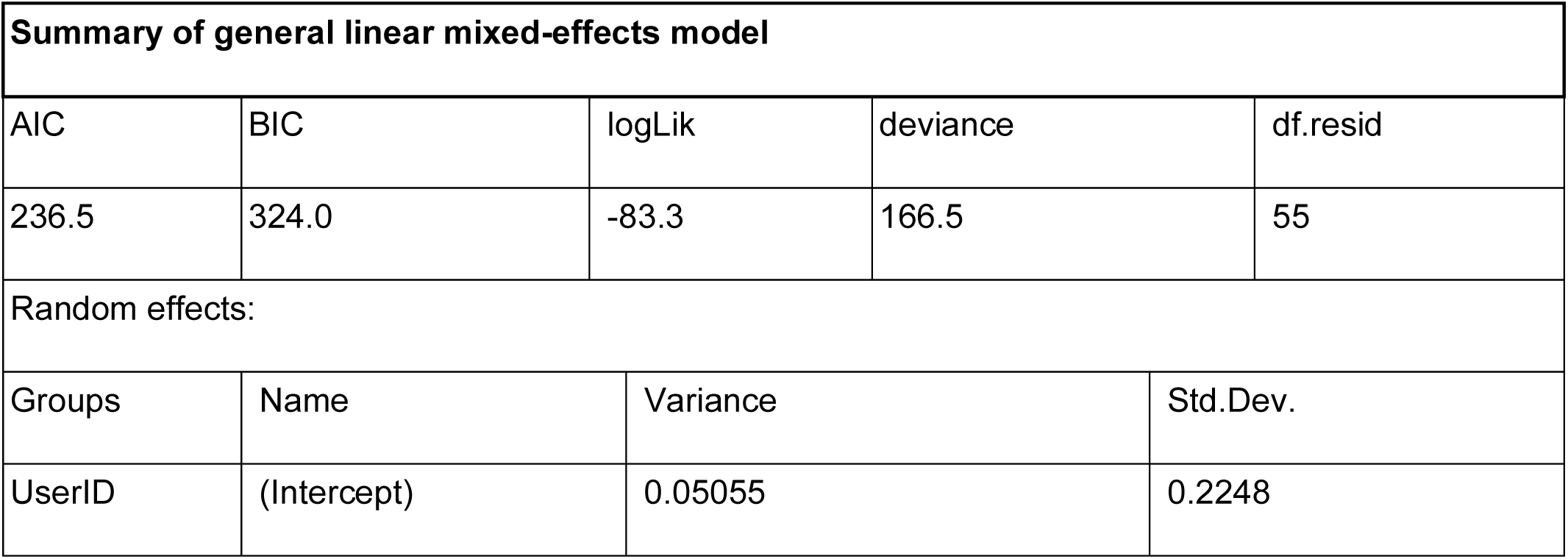
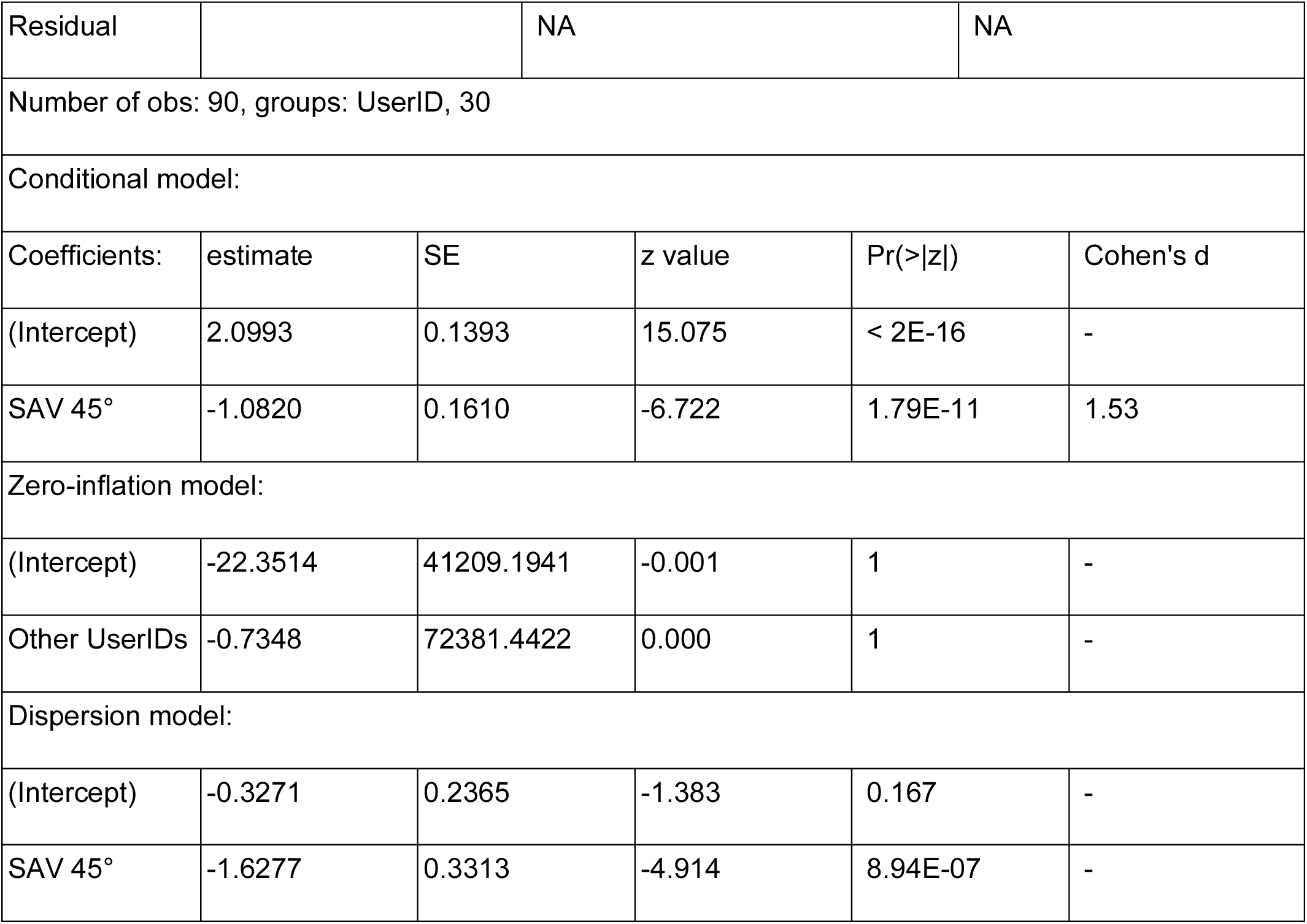

| Summary of general linear mixed-effects model |  |  |  |  |
| --- | --- | --- | --- | --- |
| AIC | BIC | logLik | deviance | df.resid |
| 236.5 | 324.0 | -83.3 | 166.5 | 55 |
| Random effects: |  |  |  |  |
| Groups | Name | Variance | Std.Dev. |  |
| UserID | (Intercept) | 0.05055 | 0.2248 |  |

|  |  |  |  |  |  |
| --- | --- | --- | --- | --- | --- |
| Residual |  | NA | NA |  |  |
| Number of obs: 90, groups: UserID, 30 |  |  |  |  |  |
| Conditional model: |  |  |  |  |  |
| Coefficients: | estimate | SE | z value | Pr(>\|z\|) | Cohen's d |
| (Intercept) | 2.0993 | 0.1393 | 15.075 | < 2E-16 | - |
| SAV 45° | -1.0820 | 0.1610 | -6.722 | 1.79E-11 | 1.53 |
| Zero-inflation model: |  |  |  |  |  |
| (Intercept) | -22.3514 | 41209.1941 | -0.001 | 1 | - |
| Other UserIDs | -0.7348 | 72381.4422 | 0.000 | 1 | - |
| Dispersion model: |  |  |  |  |  |
| (Intercept) | -0.3271 | 0.2365 | -1.383 | 0.167 | - |
| SAV 45° | -1.6277 | 0.3313 | -4.914 | 8.94E-07 | - |

- Locomotion task: Distance smoothness

Best fitting model: Generalized linear mixed-effects model with zero-inflation and dispersion correction. Distance smoothness in the locomotion task (measured as within-trial standard deviation) was modeled as the dependent variable, with viewing condition as a fixed effect. Since the design involved repeated measures, participant was included as a random effect. A zero-inflation component was specified to account for excess zero values, and dispersion was modeled as a function of the viewing condition to capture variability in response distribution.

Model assumptions: assumption violations are highlighted in red.

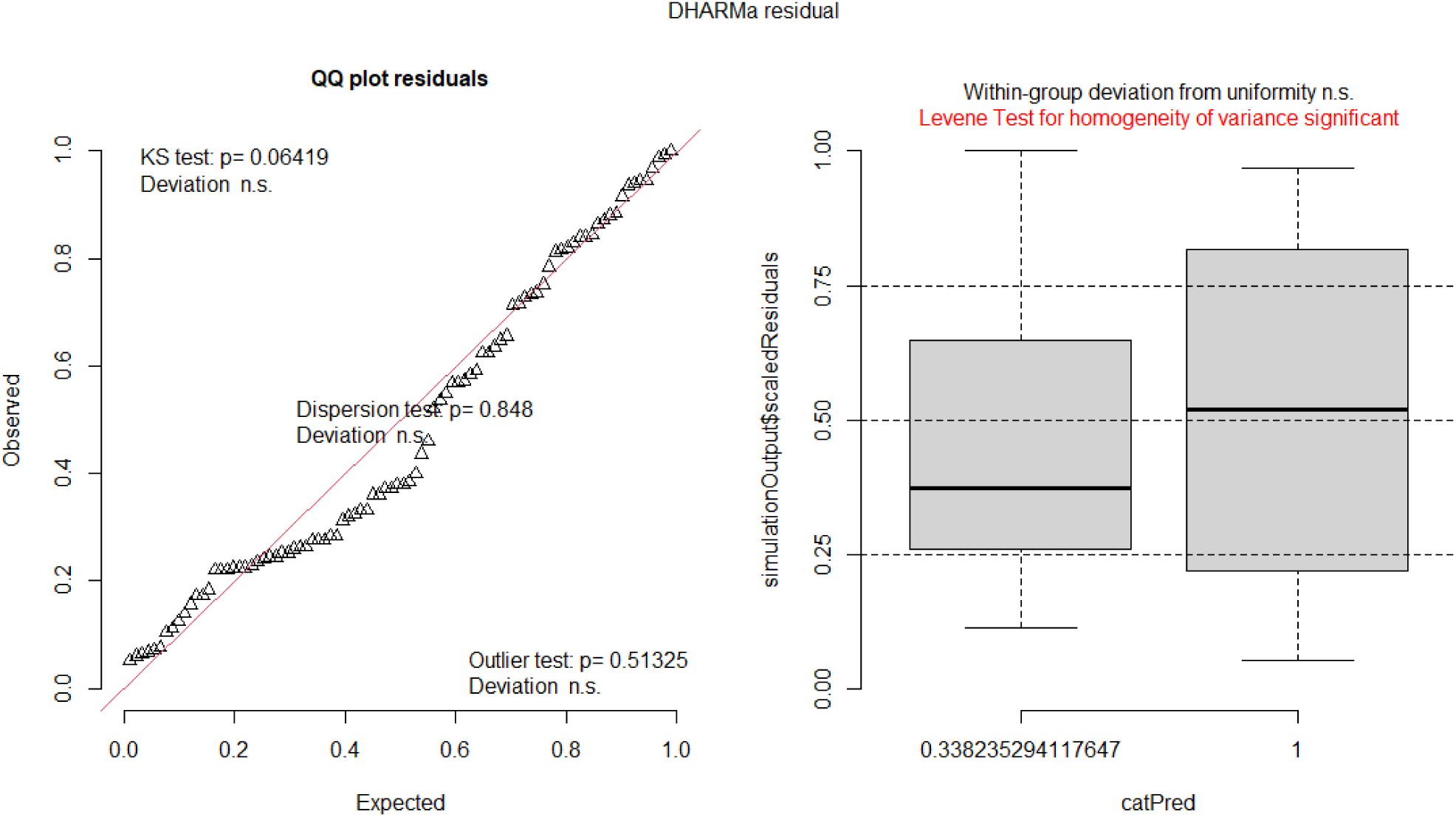

Statistical results:

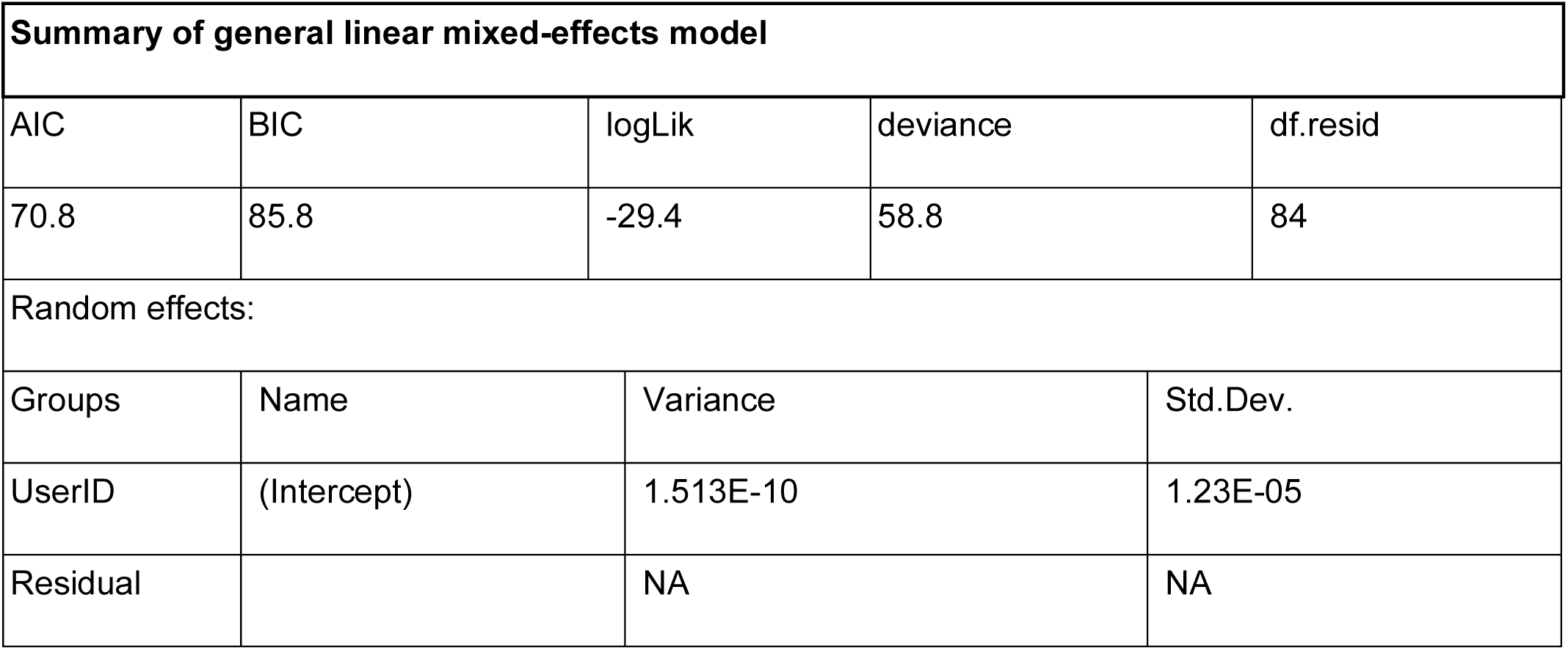
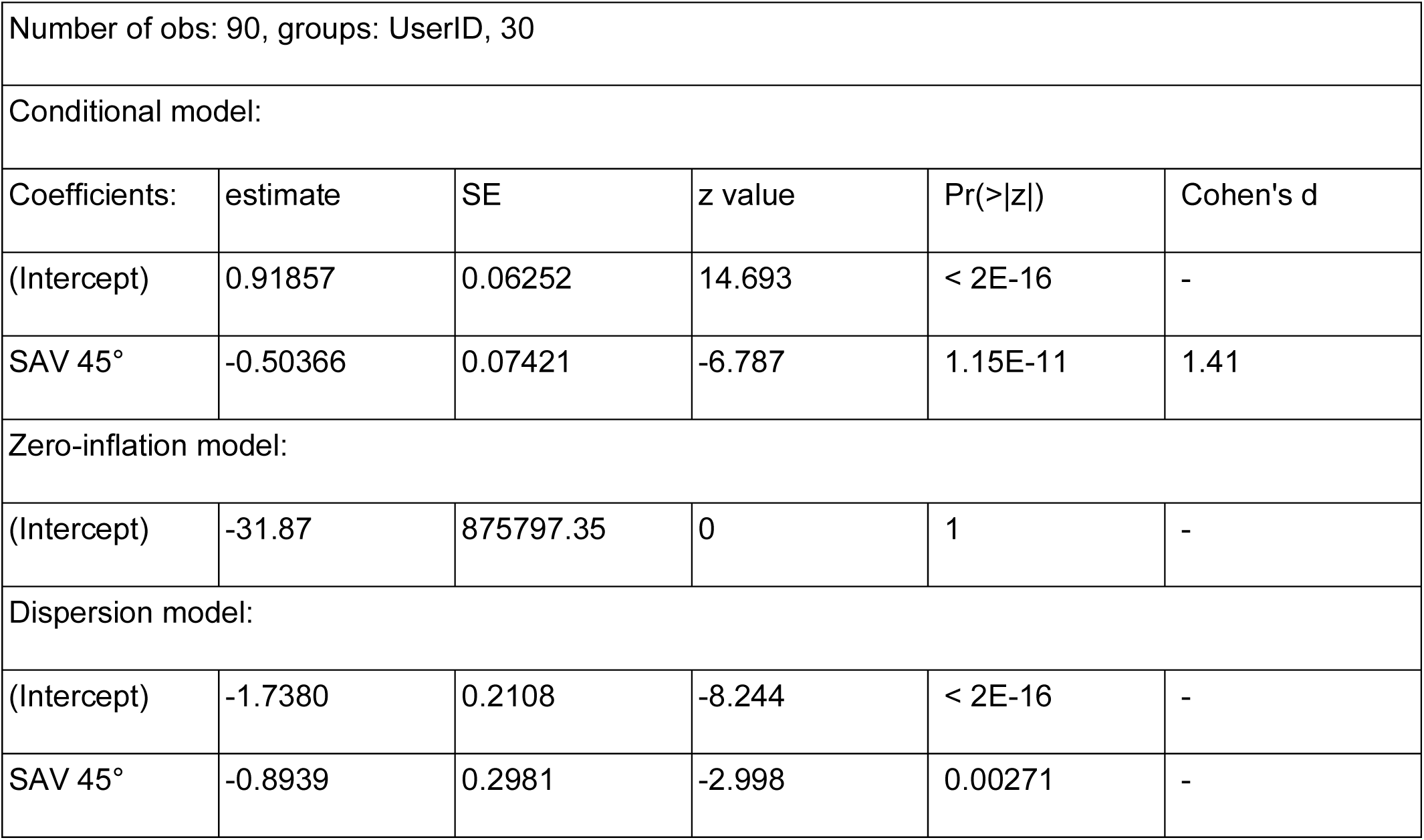

| Summary of general linear mixed-effects model |  |  |  |  |
| --- | --- | --- | --- | --- |
| AIC | BIC | logLik | deviance | df.resid |
| 70.8 | 85.8 | -29.4 | 58.8 | 84 |
| Random effects: |  |  |  |  |
| Groups | Name | Variance | Std.Dev. |  |
| UserID | (Intercept) | 1.513E-10 | 1.23E-05 |  |
| Residual |  | NA | NA |  |

|  |  |  |  |  |  |
| --- | --- | --- | --- | --- | --- |
| Number of obs: 90, groups: UserID, 30 |  |  |  |  |  |
| Conditional model: |  |  |  |  |  |
| Coefficients: | estimate | SE | z value | Pr(>\|z\|) | Cohen's d |
| (Intercept) | 0.91857 | 0.06252 | 14.693 | < 2E-16 | - |
| SAV 45° | -0.50366 | 0.07421 | -6.787 | 1.15E-11 | 1.41 |
| Zero-inflation model: |  |  |  |  |  |
| (Intercept) | -31.87 | 875797.35 | 0 | 1 | - |
| Dispersion model: |  |  |  |  |  |
| (Intercept) | -1.7380 | 0.2108 | -8.244 | < 2E-16 | - |
| SAV 45° | -0.8939 | 0.2981 | -2.998 | 0.00271 | - |

- Outdoor observation task: Identified people

Conducted test: A Mann-Whitney U test was conducted to compare the number of identified people in the outdoor observation task between the SAV 20° and SAV 45° conditions. This non-parametric test was chosen due to the non-normal distribution of the data, assessing whether there was a significant difference in median scores between the two groups.

Model assumptions:

| Shapiro test to test for normal distribution for each group |  |  |
| --- | --- | --- |
| Viewing Condition | estimate | p-value |
| SAV 20° | 0.86848 | 0.03205 |
| SAV 45° | 0.89559 | 0.08152 |
| Levene's Test for Homogeneity of Variance |  |  |
| df | estimate | p-value |
| 1 | 0.0133 | 0.9091 |
| No normality in SAV 20° group: parametric test cannot be performed |  |  |

Statistical results:

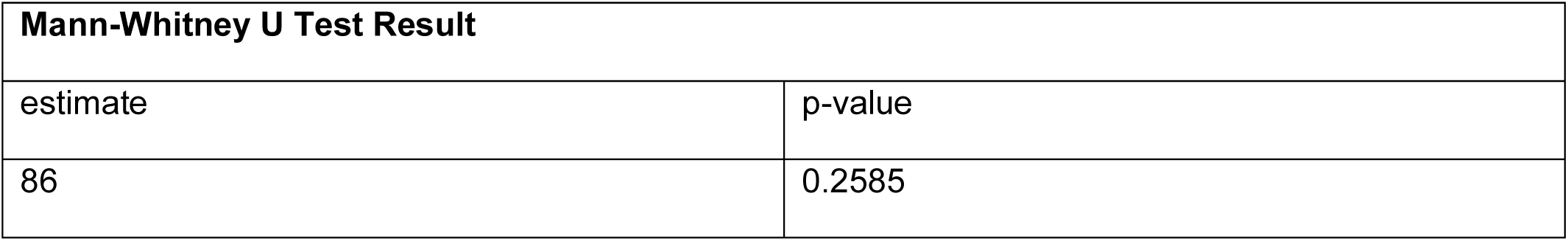

- Outdoor observation task: Attribute score

Conducted test: Since assumptions of normality and homogeneity of variance were met an independent samples t-test was conducted to compare the attribute score in the outdoor observation task between the SAV 20° and SAV 45° conditions.

Model assumptions:

| **Shapiro test to test for normal distribution for each group** |  |  |
| --- | --- | --- |
| Viewing Condition | estimate | p-value |
| SAV 20° | 0.89664 | 0.08456 |
| SAV 45° | 0.91767 | 0.1775 |
| **Levene's Test for Homogeneity of Variance** |  |  |
| df | estimate | p-value |
| 1 | 0.0133 | 0.9091 |
| Normality and homogeneity of variance: parametric test can be performed |  |  |

Statistical results:

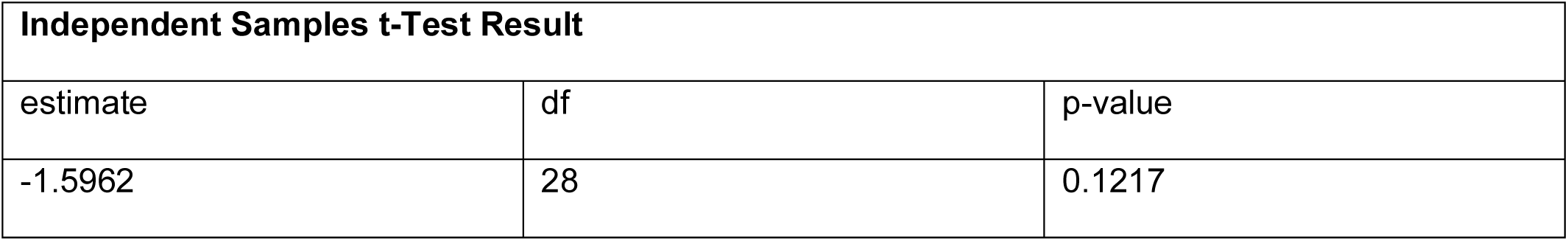

